# Natural copy number differences of tandemly repeated small nucleolar RNAs in the Prader-Willi syndrome genomic region regulate individual behavioral responses in mammals

**DOI:** 10.1101/476010

**Authors:** Maryam Keshavarz, Rebecca Krebs-Wheaton, Peter Refki, Yoland Savriama, Yi Zhang, Anja Guenther, Tanja M. Brückl, Elisabeth B. Binder, Diethard Tautz

## Abstract

The Prader-Willi Syndrome (PWS) gene region is an imprinted gene complex involved in behavioral, metabolic and osteogenic functions. We have analyzed here the variation of two families of regulatory small nucleolar RNAs (SNORD115 and SNORD116) that are coded within the PWS and are expressed from the paternal chromosome. They are organized in two tandemly repeated clusters which are naturally copy number variable between individuals. We find that the copy numbers at these loci correlate with repeatable individual test scores for anxiety that are considered to constitute a component of the “personality” of individuals. We show this for different populations and species of mice, cavies and for the anxiety component of personality tests in humans. This is also the case for an inbred mouse strain (C57Bl6) implying that copy number variation creates phenotypic variability even in an isogenic background. In transcriptome data from brain samples of this strain we find SNORD copy-number correlated regulation of target genes that are known to be involved in influencing behavior. SNORD115 has previously been suggested to regulate splicing of the serotonin receptor *Htr2c* and we confirm this in our data. For SNORD116 we provide evidence that it regulates the expression level of the chromatin regulator *Ankrd11*, which itself regulates GABA receptors, metabolic pathways, cell differentiation and osteogenesis. Intriguingly, we find that craniofacial shapes in mice correlate also with SNORD116 copy numbers. New copy number variants are generated at very high rates in mice, possibly at every generation, explaining why conventional genetic mapping could not detect this association. Our results suggest that the variable dosage of two regulatory RNAs are major determinants of individual behavioral differences and correlated traits in mammals.

## Introduction

The study of consistent individual differences in behavior has flourished over the last decades, because it has been recognised as a major contributor to differences in survival and fitness among individuals (Réale 2007; Wolf and Weissing 2012; Penke and Jokela 2016; Forkosh et al. 2019). The focus has been on traits showing relatively enduring patterns, both when measured across several occasions, as well as under different circumstances (Wilson 1998; Reale et al. 2007; Forkosh et al. 2019). This acknowledges that individuals of the same species, population, sex and even age or prior experiences show predictable differences in responding in certain ways under certain circumstances. Cornerstones of this emerging research field are to confirm that a) individuals vary from each other in behavioral responses, e.g. levels of anxiety in a specific test situation, b) this individual behavioral response shows temporal consistency and repeatability, e.g. an individual which reacts more scared then others to an Open Field Test at one occasion should react also more scared when tested at a later point in time and c) individual reactions should be consistent across different (test) situations, i.e. an individual that reacts more scared in an Open Field Test should also react more scared when confronted with an Elevated Plus Maze.

Such individual differences in behavioral patterns were found for many animal species and they resemble those found by psychologists studying human behavior, although there is an active discussion on whether they are really comparable (Uher 2011). Several studies have shown a heritable component for such behavioral differences implying some form of underlying genetic mechanisms (Dochtermann et al. 2015; Sanchez-Roige et al. 2018). Interestingly, however, such individual variation is also known to occur in genetically isogenic strains (Jakovcevski et al. 2008; Freund et al. 2013, 2015) which suggests that non-genetic mechanisms are also involved. On the other hand, a genetic locus with high rates of change between generations could potentially explain both observations.

A locus that has generally been implicated in behavioral traits in humans is the Prader–Willi syndrome (PWS) region. Mutations in this regions cause a neuro-developmental disorder which leads to several abnormalities in cognitive behaviors such as social communication, speech, anxiety, intellectual ability and decision making, but also to metabolic syndromes and craniofacial shape changes (Cassidy et al. 2012). The region is subject to imprinting, i.e. parentally biased expression of genes (Nicholls et al. 1998). The protein coding genes expressed in this region include *Ube3a*, *Snrpn* and *Magel2,* of which *Ube3a* is expressed from the maternally provided chromosome and *Snrpn* and *Magel2* from the paternally provided one. The PWS includes also several non-coding RNAs (see below). This arrangement has specifically evolved in mammals and is generally conserved among them, including humans (Zhang et al. 2014).

In mice we have previously identified the region between *Ube3a* and *Snrpn* in the PWS as one of two parentally imprinted regions that evolve particularly fast between natural populations and that could explain parentally biased mating preferences between them (Lorenc et al. 2014). Laboratory mice mutant for genes or gene regions from the PWS affect various behaviors including anxiety, activity, ultrasonic vocalization, social interaction, metabolism and foraging (Ding et al. 2008; Nakatani et al. 2009; Qi et al. 2016; Cavaille 2017; Homdli et al. 2019).

The PWS region between *Ube3a* and *Snrpn* includes two small nucleolar RNA (SNORD) gene families which are organized in large, tandemly repeated clusters known as SNORD115 and SNORD116 (Cavaille 2017) (Figure 1). The expression of both SNORD115 and SNORD116 is brain-specific and restricted to the alleles on the paternal chromosome (Ding et al. 2008).

**Figure 1:**
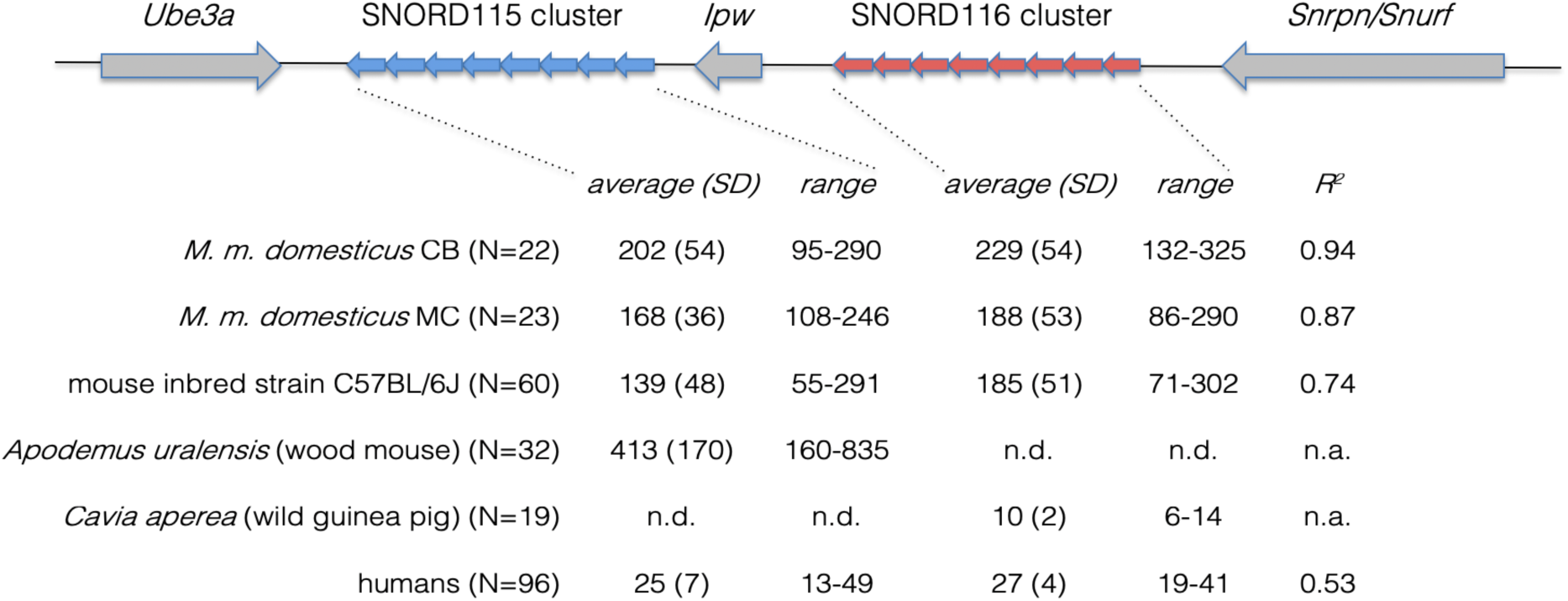
Structure of the PWS region and copy numbers of snoRNAs. The scheme depicts the genes in the cluster as arrows pointing in the direction of transcription. *Ube3a* is expressed from the maternal chromosome, the others from the paternal one. Note that the transcript structures are actually more complex and not yet fully resolved (Lorenc et al. 2014; Cavaille 2017). The tandem clusters coding for the SNORD115 and SNORD116 snoRNAs are depicted by smaller block arrows (not drawn to size). The table below provides the measures and ranges of copy numbers of the clusters in the different species analyzed in the present study. The correlation coefficients (*r*) between the numbers in the two clusters are listed to the right for the species where these are available.

SNORDs are part of a large group of small, metabolically stable snoRNAs which regulate post-transcriptional modification of their target genes (Kiss 2001; Falaleeva et al. 2017). One mechanism for snoRNA interaction is to bind via a complementary antisense region to a target RNA, whereby up to three mismatches between the antisense box and the target region can be tolerated (Kishore et al. 2010). The SNORD115 antisense element exhibits complementarity to the alternatively spliced exon V of the serotonin receptor *Htr2c* which is part of the serotonin regulatory pathway. It has been shown in cell-culture experiments and for PWS patients that there is a positive correlation between SNORD115 expression and exon part Vb usage (Kishore and Stamm 2006). However, ectopic expression of SNORD115 in non-target tissues of mice does not lead to a regulation of alternative splicing of this receptor (Raabe et al. 2019), implying that the interaction is more complex and possibly specific to the tissues of normal expression of SNORD115. Activation of the receptor by serotonin inhibits dopamine and norepinephrine release in certain areas of the brain which regulates mood, anxiety, feeding, and motoneuron functions (Stamm et al. 2017). Human SNORD116 shows a complementary sequence to an *Ankrd11* exon, allowing to suggest a direct regulation (Bazeley et al. 2008). ANKRD11 is a ankyrin repeat domain-containing protein that acts as a transcriptional co-factor with multiple possible target genes (Gallagher et al. 2015), as well as dendrite differentiation in the cerebal cortex via the BDNF/TrkB signaling pathway (Ka and Kim 2018). But this interaction has only computationally been predicted and has not yet been experimentally confirmed.

Here we ask whether natural copy number variation in SNORD115 and SNORD116 would correlate with individual differences in behavioral traits in mammals. We used standardized tests for anxiety profiles for different populations and species of mice, as well as comparable tests for wild guinea pigs and a questionnaire-based test for humans. We find that there is indeed a strong correlation between copy numbers of the respective SNORD genes and behavioral measures. Intriguingly, this is not only found for wildtype strains, but also for the common laboratory mouse inbred strain C57BL/6J. Using transcriptomic analyses, we show that the predicted regulation of *Htr2c* and *Ankrd11* can indeed be observed and that the network of genes affected by the copy number variation of SNORD116 can explain the behavioral and osteogenic phenotypes. Our data suggest further that new alleles with different copy numbers are generated at an exceptionally high rate. Hence, we conclude that this variation could be a basis for the long sought molecular mechanism for the high variance of individual behavioral traits in families and populations of mammals.

## Results

### ddPCR to measure repeat numbers

Given the possible molecular links to the regulation of behavioral responses, we were interested to measure copy number variation within the SNORD115 and SNORD116 clusters. Both snoRNAs are part of a larger transcription unit, from which they are processed (Cavaille 2017). In the mouse reference genome, the repeat unit length for SNORD115 is 1.97kb including the RNA coding region with a length of 79nt. For SNORD116 the repeat unit length is 2.54kb and the RNA has a length of 94nt. Not all copies are annotated in the reference genome, but in the genome sequence itself, SNORD115 is represented with 143 copies and SNORD116 with 70 copies, with a 50kb sequence gap (S1 Figure).

Typing copy numbers in this range has long been a technological challenge, explaining also why natural copy number variation in this region has not received much attention so far. The classic quantitative PCR approaches do not have enough power to distinguish e.g. 150 from 160 copies. Also long read sequencing procedures (e.g. PacBio or Oxford Nanopore) come to their limits in this region, since they stretch across several hundred kb, which is beyond the standard limits of these technologies. S2 Figure shows an example of the non-suitable read coverage that is obtained with these technologies in the region spanning *Ube3a*-*Snrpn*. But a new technology, droplet digital PCR (ddPCR), has been developed to achieve the necessary resolution (Pinheiro et al. 2012; Quan et al. 2018). We have previously validated this technology by comparing predicted copy numbers based on sequencing read depth from whole mouse genome sequences with measured copy numbers based on ddPCR assays (Pezer et al. 2015) S3 Figure shows that there is a very good correspondence between ddPCR results and read depth calculation for different CNV loci with high copy numbers. A possibly confounding factor for ddPCR would be that primers may not bind equally in all repeat units, due to polymorphisms in the primer binding sequence. To reduce the chance of polymorphism in the primer binding sites, we placed them inside the SNORD RNA coding parts. Still, a survey of all described SNORD variants suggested that some polymorphism exists also in these regions. Hence, we have tested two different sets of primers and found that both yield highly congruent results (see Methods). We have therefore conducted the bulk of the experiments with only one set of primers.

To control the technical replicability, we have run the ddPCR reactions for the individuals in triplicates. These show a very high correspondence, with average standard deviations of 3.54 for SNORD115 and 3.15 for SNORD116 in the *Mus* samples (S1 Table). We tested also biological replicates by using ear samples and brain samples from the same individuals. We found also highly congruent results in these tests (S2 Table), implying also that the copy number in ear samples are a very good proxy for brain samples.

In contrast to the long-range sequencing approaches discussed above, the optical mapping procedure developed by Bionano has at least the chance to generate information for fragments longer than 500kb. But it has also its limitations. First, it is restricted to regions where the repeat units include at least one recognition site for the enzyme used for labelling. In the mouse this applies only to the SNORD116 repeat units, which have one recognition site for Nt.BspQI and two sites for Dle1 (these are the two alternative enzymes that are used for labelling). Second, the procedure requires to obtain a substantial amount of high-quality DNA and it is complex and expensive. Still, we have obtained Bionano optical mapping data for seven individuals. Unfortunately, even with this technology and with 100x coverage of the whole genome, we found that reads spanning the whole SNORD116 cluster are very rare. In the end we could get enough suitable fragments to distinguish two alleles for one of the animals only (S4 Figure). For this animal we were able to confirm that the SNORD116 region is indeed a contiguous tandem repeat (i.e. the gap in the reference sequence could be filled with repeat units). Further, we applied ddPCR to this animal and we found that the manually counted number of copies from the optical map (287) matched very closely the value that we obtained with ddPCR (293) (S4 Figure), providing another proof for the validity to use ddPCR. Because of the rareness of full length fragments spanning the full repeat regions and the high costs, optical mapping is not suitable for typing a large number of animals, especially not when the repeat units do not include the recognition sites for labelling (as it is the case for the SNORD115 repeats in mouse and both types of SNORD repeats in humans). Hence, ddPCR is currently the only method that has shown a sufficient power and reproducibility to quantify the copy numbers in tandem repeat regions, which is the key to the results described below (see also discussion).

### SNORD copy number variation

Using the ddPCR assay, we tested first individuals derived from two populations of *M. m. domesticus* that were originally caught in the wild in Germany (CB) and France (MC) and which were kept under outbreeding conditions (Harr et al. 2016). We find an average of 202 SNORD115 copies in CB animals and 170 in MC. A similar difference exists for SNORD116 copies, with 229 in CB and 188 in MC (Figure 1). Although there is a large variation within the populations, these differences between the populations are significant (t-test; p=0.03 for SNORD115 and p=0.015 for SNORD116; all data normally distributed, Shapiro-Wilk normality test p>0.21). Intriguingly, we observe also a strong co-variation of copy number between the two SNORD clusters in both populations, with highly significant correlation coefficients *r* (Figure 1). Such a co-variation is unexpected, since the repeat units are separated by a non-repeated region and since they share no sequence similarities. To test whether a possibly unrecognized technical factor leads to this correlation, we typed also the two other CNV loci in the mouse genome that show particularly high copy-number range (*Sfi1* and *Cwc22* – see S3 Figure) for the same set of individuals. *Sfi1* has between 18-98 copies in the CB population and 18-121 copies in the MC population, *Cwc22* shows high copy number variation in the CB population only (18-99 copies) (Pezer et al. 2015). We find that neither of these two loci shows a correlation with SNORD115 or SNORD116 copy numbers (S3 Table). As a further control for the validity of the correlation between SNORD115 and SNORD116 copy numbers, we compared read coverage from the whole genome sequence data set of *Mus musculus* populations included in Harr et al. (Harr et al. 2016). We find indeed also a high correlation (*r*^2^ = 0.64) between read coverage for SNORD115 and SNORD116 in individuals (S5 Figure), supporting the observations from the ddPCR data.

### Copy numbers and RNA expression

Given the proven efficiency of ddPCR for copy-number determination, we used it also to quantify RNA expression (Quan et al. 2018) in individuals from the wildtype population. The measured SNORD gene copy numbers show a highly significant regression with the expression of the respective SNORD RNAs in the brain (Figure 2A, B) (S4 Table), suggesting a direct relationship between copy number and expression level.

**Figure 2:**
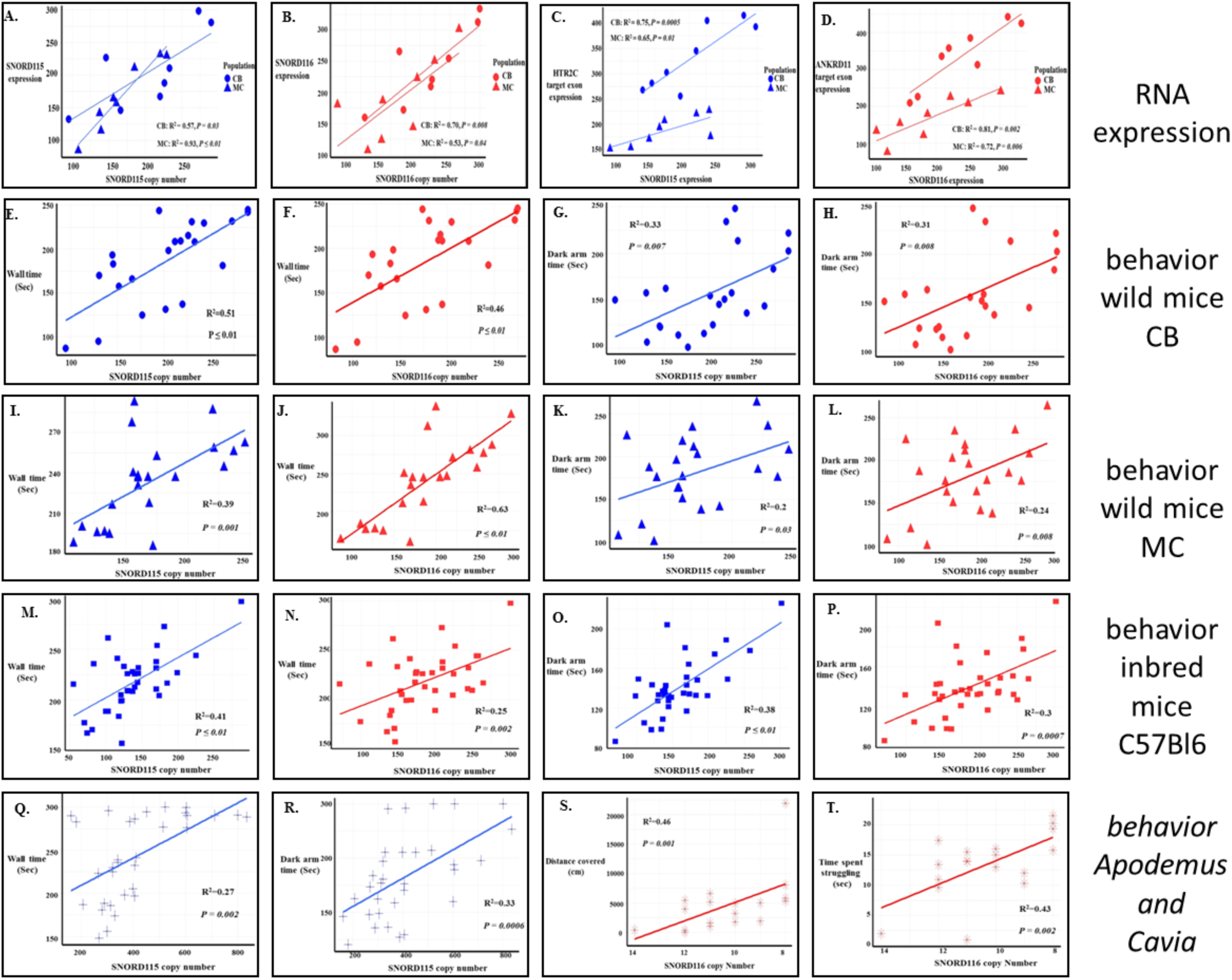
Regression between SNORD115 (blue) and SNORD116 (red) copy numbers and expression, anxiety-like behavior and skull shape. A) and B) Regression with the respective RNA expression values. C) and D) Regression between the respective SNORD expression and the splice-site regulated target RNAs. All values were determined by ddPCR in whole brain RNA preparations from eight outbred animals each of the CB and MC populations. The y-axis shows the RNA molecule copy number per µL input (see Methods) after normalization. β-catenin was used to normalize the data for HTR2C and ANKRD11 expression and SNORD66 was used to normalize the data from SNORD115/116 expression. The animals constitute a subset of the 45 animals used for the overall copy number variation reported in Figure 1 and were chosen to reflect the spread of the variation. E, F, I, J, M, N and Q show regression analysis between Time at wall in the Open Field test and SNORD copy number in wild mice CB (E, F), wild mice MC (I, J), inbred mice C57Bl6/J (M, N) and *Apodemus uralensis* (Q). G, H, K, L, O, P and R show regression analysis between Dark arm time in the Elevated Plus Maze test and SNORD copy number in CB (G, H), MC (K, L), inbred mice C57Bl6/J (O, P) and *Apodemus uralensis* (R). S,T: Regression analysis between SNORD116 copy number and distance covered (cm) from the Open Field (S) and time spent struggling from the Struggle Docility tests (T) in the wild guinea pig *Cavia aperea*. Note that the behavioral tests for cavies differ from the tests for the other species. Lower scores represent higher anxiety, i.e. the X-axis is reversed for better comparability. In all tested rodent species in this study, animals with higher copy numbers have a higher relative anxiety score.

Further, we asked whether the target RNA expressions correlate with their respective SNORD expression. The target of SNORD115 is the alternatively spliced exon V of *Htr2cr* (Kishore and Stamm 2006), which is split into two halves through the alternative splice – exonVa and exonVb (S6 Figure). We used primers that span specifically the exonVb to exonVI junction and we find a significant positive correlation for this variant with SNORD115 copy numbers in both wildtype populations (Figure 2C). The predicted target of SNORD116 is exon IX of *Ankrd11* (Bazeley et al. 2008) (S6 Figure). Since no alternative splice was described for this so far, we placed the primers close to the predicted SNORD116 binding site and we find a significant correlation with SNORD116 copy numbers (Figure 2 D) (all data in S4 Table). Note that these patterns are confirmed and extended by the RNASeq analysis described below.

### Behavioral tests

To assess whether the SNORD gene copy number variation correlates also with behavioral scores of the mice, we used two standard tests in mice, the Open Field Test and the Elevated Plus Maze test. Both measure anxiety profiles and have been used before in studies of individual behavior (Jakovcevski et al. 2008; Freund et al. 2013). We determined which measurement scores generated in these tests showed significant repeatability for individuals over the course of the experiment, using intra-class correlation coefficients (see details in Methods). These pre-tests for repeatability were carried out on individuals from the MC population, the C57Bl6/J strain and the wood mouse *Apodemus,* three times each with 4-week intervals between the tests (S5 Table, S6 Table). Among the seven measurements taken, only one was highly repeatable for each test, namely “Time at wall” for the Open Field Test and “Dark arm time” for the Elevated Plus Maze test (Table 1). Hence, we used these two measurements for all mouse experiments, both as the first-time measurement for each individual, as well as in a mixed model analysis.

**Table 1:**
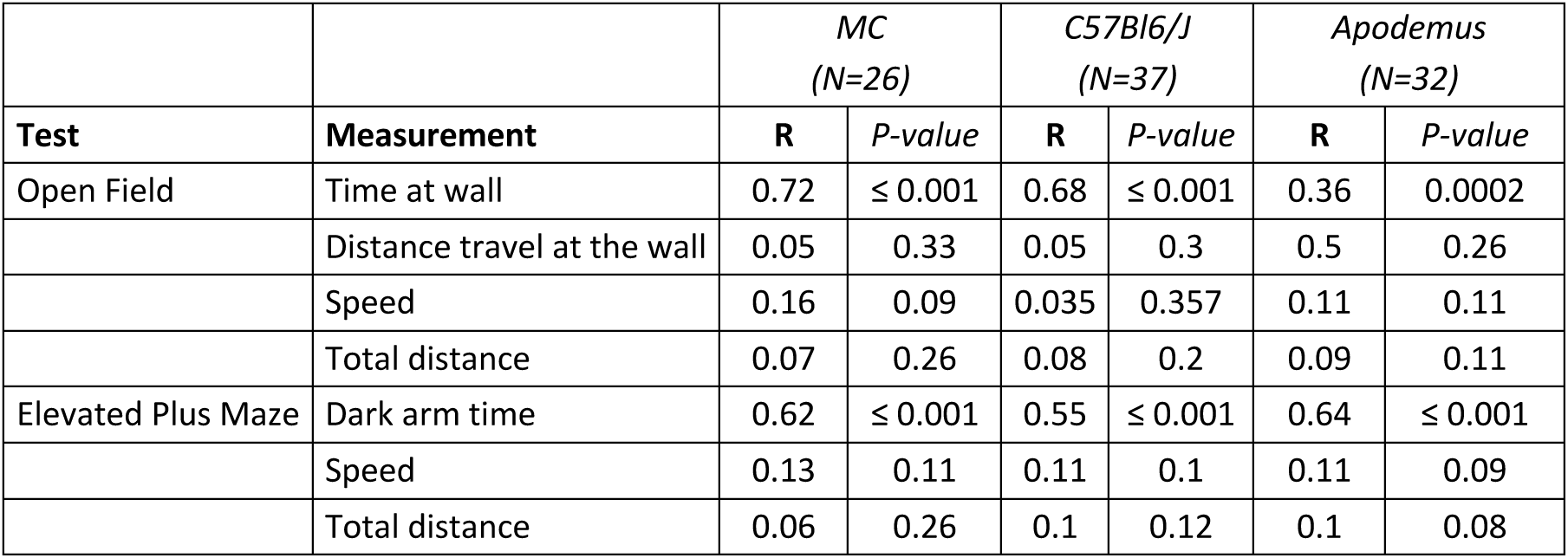
Repeatability (**R**) for behavioral tests.

| Test | Measurement | MC<br>(N=26) |  | C57Bl6/J<br>(N=37) |  | Apodemus<br>(N=32) |  |
| --- | --- | --- | --- | --- | --- | --- | --- |
|  |  | R | P-value | R | P-value | R | P-value |
| Open Field | Time at wall | 0.72 | ≤ 0.001 | 0.68 | ≤ 0.001 | 0.36 | 0.0002 |
|  | Distance travel at the wall | 0.05 | 0.33 | 0.05 | 0.3 | 0.5 | 0.26 |
|  | Speed | 0.16 | 0.09 | 0.035 | 0.357 | 0.11 | 0.11 |
|  | Total distance | 0.07 | 0.26 | 0.08 | 0.2 | 0.09 | 0.11 |
| Elevated Plus Maze | Dark arm time | 0.62 | ≤ 0.001 | 0.55 | ≤ 0.001 | 0.64 | ≤ 0.001 |
|  | Speed | 0.13 | 0.11 | 0.11 | 0.1 | 0.11 | 0.09 |
|  | Total distance | 0.06 | 0.26 | 0.1 | 0.12 | 0.1 | 0.08 |

### Behavioral scores and copy number

Using the first-time measurements for each individual, we find a highly significant regression between SNORD copy numbers and test scores in both the CB (Figure 2 E-H) and the MC (Figure 2 I-L) wild house mouse populations (data in S5 Table). The results from both populations show that animals with higher copy numbers have higher relative anxiety scores.

For the repeated measurements per individual, we constructed univariate mixed models, including animal ID as random effect and triplicate measurements of SNORD copy number from ddPCR as fixed effects. This analysis was done to account for the potential differences over the course of the behavioral experiments (three times measurement) as well as technical error from triplicate measurements from ddPCR for SNORDs copy number calculation. Regression analyses of SNORD copy number variation and behavioral variables confirmed the positive significant regression in both CB (in Open Field Test for SNORD115: R^2^=0.3, P = 0.001; SNORD116: R^2^ = 0.27, P = 0.001 and in Elevated Plus Maze for SNORD115: R^2^=0.22, P = 0.004; SNORD116: R^2^ = 0.20, P = 0.006) and MC (in Open Field Test for SNORD115: R^2^=0.24, P = 0.001; SNORD116: R^2^ = 0.41, P = 0.0001 and in Elevated Plus Maze for SNORD115: R^2^=0.23, P = 0.003; SNORD116: R^2^ = 0.22, P = 0.006).

To control whether the two other known high copy CNV loci would also show correlation with the behavioral scores, we analyzed the population samples for *Sfi1* and *Cwc22* copy numbers (see above). We find that both loci show no significant correlation with the behavioral scores in the populations tested (S3 Table), i.e. the correlations exist only for SNORD115 and SNORD116.

### Inheritance pattern

Given the high variability of SNORD gene copy numbers in the mouse populations, one can ask whether they are subject to frequent changes, such as unequal cross-over events during meiosis. Given the total length of the repeat regions, it is difficult to test this directly, since the haplotypes cannot be distinguished by most conventional techniques (see above). However, we approached this question by comparing the inheritance of copy numbers in nine families of the MC population. We find that the offspring show a large variation, exceeding the spread of numbers measured for the parents in 7 out of the 9 families (Figure 3). Most importantly, if the alleles in the offspring would be only combinations of the parental alleles, one should expect at most four allele classes in the offspring. But two families with more than four offspring (families 1 and 7) show additional allele classes (even when one takes the standard deviation of measurement errors into account: S7 Table), suggesting that new length alleles have been generated in a single generation.

**Figure 3:**
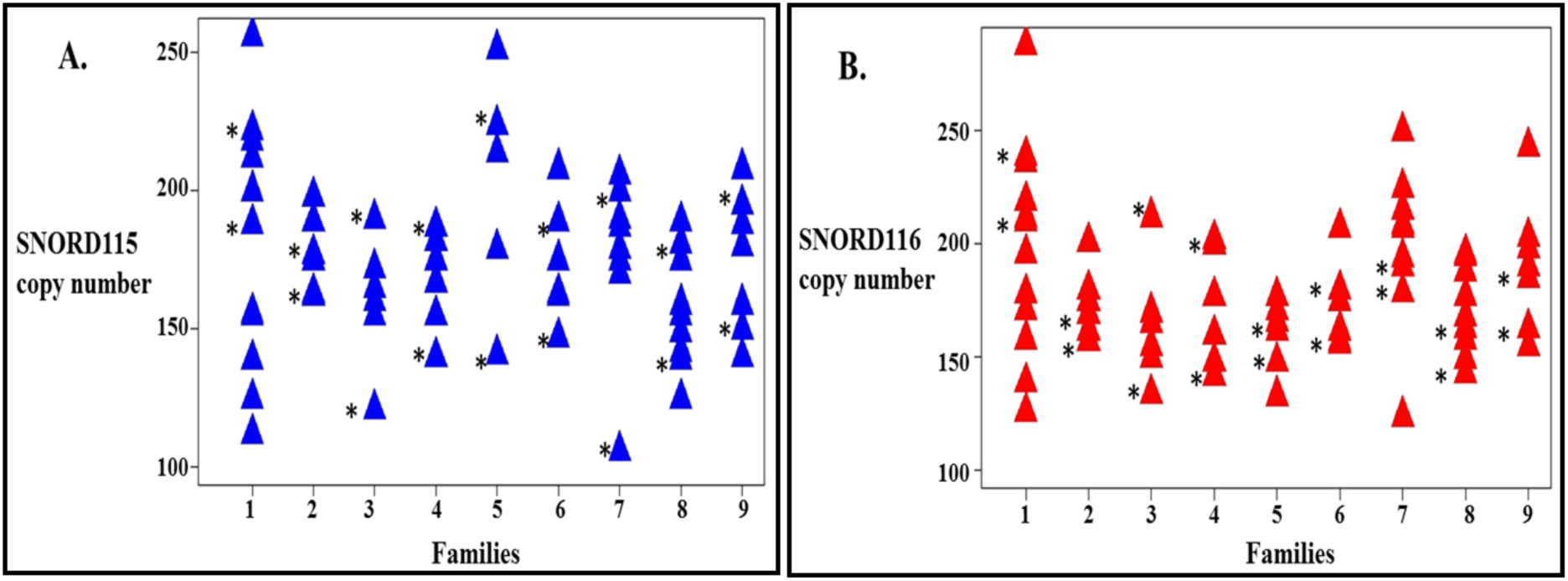
Inheritance patterns of SNORD gene copy numbers (CN) in families. Nine families of the MC population were tested and the copy numbers were determined for each individual (blue triangles for SNORD115 and red for SNORD116). The parental copy numbers are marked with a star.

The experiment shows also that new large variation is generated in each generation, in line with the general observation that anxiety traits tend to differ between parents and offspring, as well as among the offspring. To confirm that this is also the case here, we subjected 26 individuals of the offspring to the behavioral tests described above. We find indeed a large variation in behavioral scores, but also the expected significant regression with copy numbers (Open Field Test for SNORD115: R^2^=0.55, P << 0.001; SNORD116: R^2^ = 0.40, P = 0.004; Elevated Plus Maze for SNORD115: R^2^=0.33, P=0.002; SNORD116: R^2^ = 0.31, P = 0.004) (data in S5 Table). The univariate mixed-model regression analysis also showed a positive significant regression between SNORD copy number and behavioral scores (Open Field Test for SNORD115: R^2^=0.34, P << 0.001; SNORD116: R^2^ = 0.3, P << 0.001; Elevated Plus Maze for SNORD115: R^2^=0.24, P=0.003; SNORD116: R^2^ = 0.22, P = 0.003).

### Inbred strain variation

It has been shown previously that anxiety scores can differ even between individuals in inbred strains (Jakovcevski et al. 2008; Freund et al. 2013). Hence, we asked whether inbred mice would also show copy number variation for SNORD115 and SNORD116. We used the C57BL/6J mouse strain for this purpose, which is known to harbor extremely little variation in the form of SNP polymorphisms (Zurita et al. 2011). We genotyped 60 individuals of both sexes (30 female and 30 male). We find that there is indeed a large variation of copy numbers in these inbred mice, comparable to the spread seen in the outbred cohorts (Figure 1) (S1 Table). 20 mice of each sex were then subjected to the behavioral tests. We found no significant difference between males and females in these tests (ANOVA; p=0.62 for SNORD115 and p=0.28 for SNORD116) but we could confirm the strong correlation with copy numbers (Figure 2 M-P). Similarly, the univariate mixed-model regression analysis also showed significant regression between SNORD copy number and behavioral scores (Open Field Test for SNORD115: R^2^=0.40, P << 0.001; SNORD116: R^2^ = 0.32, P = << 0.001; Elevated Plus Maze for SNORD115: R^2^=0.33, P << 0.001; SNORD116: R^2^ = 0.3, P = << 0.001).

### Target gene analysis from transcriptome data

To investigate whole transcriptome changes in response to copy number variation of the SNORD genes, we choose ten C57BL/6J individuals from the previous experiment, representing a spread of copy numbers from low to high and sequenced RNA from their brains. First, we sought to confirm that the target genes that we had tested in the wildtype background by ddPCR (see above) would show the same patterns in the RNASeq data from the inbred strain.

The first analysis focused on *Htr2c* as target of SNORD115. The expected alternative splice is in exon V, which results in a 95nt long exon part (exon Vb) that is specific for one splice variants, while the rest of the mRNA is shared between both (S6 Figure). We have therefore determined RNASeq read coverage for the exon Vb part as well as exon VI and compared each to SNORD115 copy numbers from the respective animals. We find a significant positive correlation for exon Vb, but none for exon VI (R^2^ = 0.43 vs -0.12; P = 0.04 vs. 0.3) (S8 Table). This corroborates the original finding by (Kishore and Stamm 2006) that increasing the dose of SNORD115 transcripts in cell culture experiments leads to an increase in the relative concentration of exonVb into the full transcript. It confirms also our results from the ddPCR experiments in the wild type populations reported above. We have further also analyzed the edited sites in exon Vb in our RNASeq data. We find the same sites that were previously described (Burns et al. 1997; Kishore and Stamm 2006) (S6 Figure), but we do not find the additional edited sites that were reported in the ectopic SNORD115 expression study (Raabe et al. 2019). Given that this study had also not found the alternative splicing signal, it would seem that it is not comparable with the normal expression of SNORD115.

Computational analysis has predicted *TAF1*, *RALGPS1*, *PBRM1, CRHR1,* and *DPM2* as five additional possible targets for SNORD115 (Kishore et al. 2010). However, our RNAseq data analysis showed no significant correlation between SNORD115 copy number with the expression of these five genes, only a tendency for a negative correlation for *DPM2* (R^2^ = 0.32, P = 0.09) (S8 Table).

For *Ankrd11* we correlated read coverage for the exon IX 5’-part including the PCR fragment that we had used for the ddPCR study above, as well as the SNORD116 binding site (S6 Figure). This shows a significant positive correlation to SNORD116 copy number (R^2^ = 0.5; P = 0.02) (S8 Table). We have also analyzed read coverage of the exons preceding exon IX to explore possible alternative splicing. Interestingly, exon VIII shows no significant correlation with SNORD116 copy number (R^2^ = 0.009; P = 0.79), while exon VII shows a strongly negative one (R^2^ = −0.53; P = 0.01) (S8 Table). This suggests that alternative splicing happens in this region, but the current data did not allow us to resolve this unequivocally. However, in addition to the several annotated splice variants (compare S6 Figure), we find further ones in the RNASeq data that will require a much deeper analysis. Also, in contrast to *Htr2C*, we find no evidence for RNA editing around the predicted SNORD116 binding site in *Ankrd11*. Hence, we conclude that the mechanism for regulation may be different, but this is not unexpected given the diversity of regulatory interactions that are known for C/D-box snoRNAs (Falaleeva et al. 2017).

### Whole transcriptome analysis

Given that we test a range of copy number variants, rather than two states, it is not possible to do a conventional analysis of significantly differentially expressed genes across the whole genome, where only two states are compared. Hence, we used correlation coefficients (*R^2^*) as a basis for identifying possible target genes. Since SNORD116 appears to regulate *Ankrd11* and since ANKRD11 is a chromatin regulator which influences the expression of downstream genes by binding chromatin modifying enzymes like histone deacetylases, we started with a list of previously identified target genes of ANKRD11 (Gallagher et al. 2015). We were able to match the data for 635 predicted target genes from the (Gallagher et al. 2015) study with our data from the whole transcriptome analysis. We used these to correlate the expression data of these genes from our whole transcriptome analysis to both the *Ankrd11*-exon IX 5’-part ddPCR and the SNORD116 copy numbers. When applying a cutoff of *R^2^*=0.4 for both correlations, we find 72 positively correlated and 32 negatively correlated genes (S9 Table). A subunit of the GABA A receptor (*Gabrg1*) shows the strongest positive correlation among the known target genes. *Gabrg1* is known to play a crucial role in the modulation of the anxiety response in both humans and mice (Nuss 2015).

In a second step, we applied the same correlation analysis to all genes in the transcriptome, using a correlation with either SNORD115 or SNORD116 copy numbers. We found an additional 67 candidate genes for SNORD115 and 89 for SNORD116 (S9 Table). This suggests that the transcriptome effects of the two SNORD genes go beyond the known target pathways, although this could of course be a secondary effect. Note that the *Ankrd11* level correlates as a whole with SNORD116 copy number, as predicted from the ddPCR on the *Ankrd11*-exon IX 5’-part results above, i.e. the exon which represents most of the full-length RNA (S6 Figure). In contrast, the level of the whole *Htr2c* transcript does not correlate with SNORD115 copy number, since the alternative splice changes are only a minor part of it (S6 Figure).

### SNORD116 copy number and craniofacial features

Given that a previous study of a particular mutation in the *Ankrd11* gene has shown craniofacial abnormalities (Barbaric et al. 2008), we tested whether SNORD116 copy number variation would also correlate with craniofacial features in the C57BL/6J mouse strain. We used a 3D landmarking approach to quantify differences in skull shapes based on CT scans, following the procedures described in (Pallares et al. 2015). A multivariate regression of the shape data (Procrustes coordinates) on SNORD116 copy number indicated a strong relationship (percentage predicted = 7.97%; p-value < 0.0001; 10,000 rounds of permutation) (Figure 4A).

**Figure 4:**
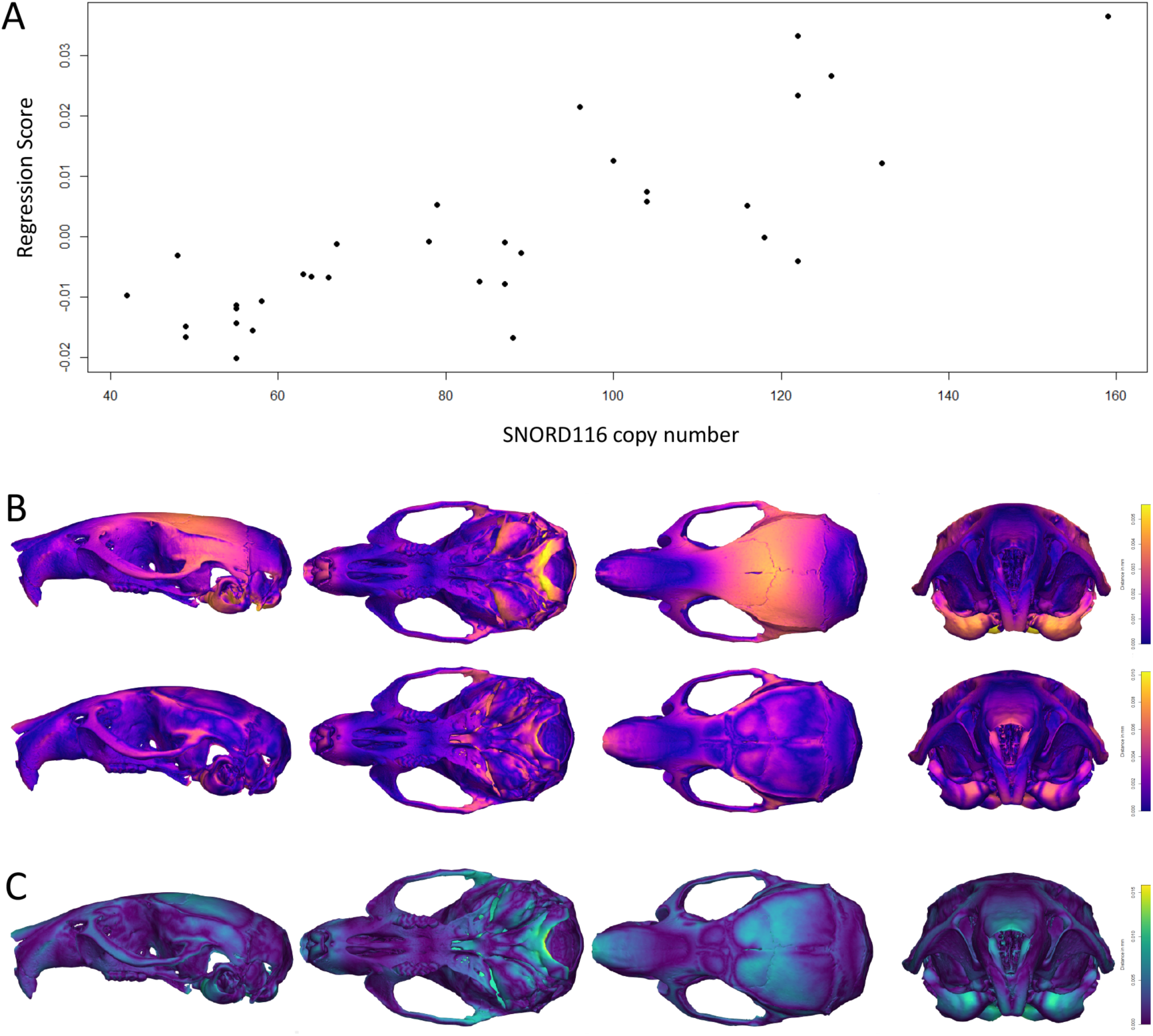
Morphometric analysis of mouse (C57Bl6) crania in dependence to SNORD116 copy number variation. A) Shape scores obtained from the multivariate regression plotted against SNORD116 copy number. B) Shape change associated with a low SNORD116 copy number (top) and high copy number (bottom) visualized by warped 3D surfaces of mouse skulls representing the deviation of each predicted shape and the overall mean shape (warm colors indicate high deviation while cold colors stand for low deviation measured in Procrustes distance). C) Same as B, but differences between predicted shape change for a low and high SNORD116 copy number.

The variation in SNORD116 copy number is associated with skull shape changes mostly localized at the back of the skull (i.e. braincase, squamosal, and occipital) with a low copy number being linked to more slender and straight skulls while a high copy number is related to more rounded and bent skulls with an enlarged braincase (Figure 4B, C).

### Other rodents

Given that the arrangement of the PWS region with the two clusters of snoRNAs is conserved throughout mammals (Sato 2017), we were interested to assess in how far one can trace the association between copy number variation and anxiety scores also in other species.

We have first analyzed a second mouse species, the wood mouse *Apodemus uralensis*, which is separated from *Mus musculus* since about 10 million years and which has different ecological adaptations. Still, given its general similarity to house mice, we applied the same phenotyping scheme and verified that the phenotype scores are repeatable for individuals (Table 1). From the currently available genomic data, we could only retrieve sufficient information for the SNORD115 cluster, but this has even higher copy numbers than in *Mus musculus* (Figure 1). We find indeed a significant regression with SNORD115 copy numbers and behavioral scores (Time at wall from Open Field and Dark arm time from Elevated Plus Maze) in *Apodemus* as well (Figure 2 Q, R). Mixed-model regression analysis also showed a significant regression between SNORD115 copy number and behavioral measurements (in Open Field Test for SNORD115: R^2^=0.35, P << 0.001; and in Elevated Plus Maze for SNORD115: R^2^=0.24, P = 0.001) (all data in S6 Table).

As a second, even more distant rodent, we used a cavy species, *Cavia aparea,* the wild congener of domesticated guinea pigs. For this species we used two previously established behavioral tests that can be considered to reflect anxiety behavior and were shown to be repeatable (Guenther and Trillmich 2015), but differ in their execution and their scores from the mouse tests (see Methods). The current genome sequence of a close relative, *Cavia aperea* f. *porcellus,* provides only the annotation for SNORD116 repeats, hence we focused our analysis on this cluster. We find a much smaller range of copy numbers than in mouse (Figure 1), but still a significant regression with the specific cavy anxiety scores from the two tests (Figure 2 S,T) (all data in S10 Table). As with the other rodents in this study, individuals with higher copy numbers showed higher anxiety.

### Humans

To assess a possible association between SNORD copy numbers and personality traits in humans, we tested a subset from a cohort of 541 healthy individuals that had taken part in a study based on the Tridimensional Personality Questionnaire (TPQ) which is designed to explore novelty seeking, harm avoidance and reward dependence (Cloninger et al. 1991; Weyers et al. 1995). From these individuals, we chose the top 48 each for low and high anxiety scores (see Methods) and typed them for SNORD copy numbers. SNORD115 in humans is represented by a single class only, while the SNORD116 family is split in three subclasses, for which we designed different ddPCR assays. We find significant differences of copy numbers between the two groups (Table 2). Particularly significant are SNORD115 and SNORD116_2. The latter is the variant that is predicted to bind to *Ankrd11* exon IX, while the possible target genes for the other two SNORD116 variants are not yet clear. These latter ones show also generally only little copy number variation (Table 2) (all data in S11 Table). However, we note that the direction of the correlation is different between rodents and humans. In humans, the relatively higher anxiety group has the smaller number of copies, while it is the other way around in the three tested rodent species (see above).

**Table 2:** SNORD copy numbers (mean) for humans scoring low or high on anxiety and activity

| ddPCR assay | SNORD115 | SNORD116_1 | SNORD116_2 | SNORD116_3 | all SNORD |
| --- | --- | --- | --- | --- | --- |
| low anxiety and activity group, N=48 (SD) | 28 (8.0) | 6 (1.1) | 16 (4.4) | 8 (1.4) | 57 (12.6) |
| high anxiety and activity group, N=48 (SD) | 21 (4.5) | 6 (0.8) | 12 (1.7) | 7 (1.1) | 47 (6.8) |
| p-value<sup>#</sup> | $7.2 \times 10^{-6}$ | 0.29 | $1.1 \times 10^{-5}$ | 0.06 | $4.4 \times 10^{-6}$ |

<sup>#</sup> unpaired t-test, two-tailed

**Table 3.**
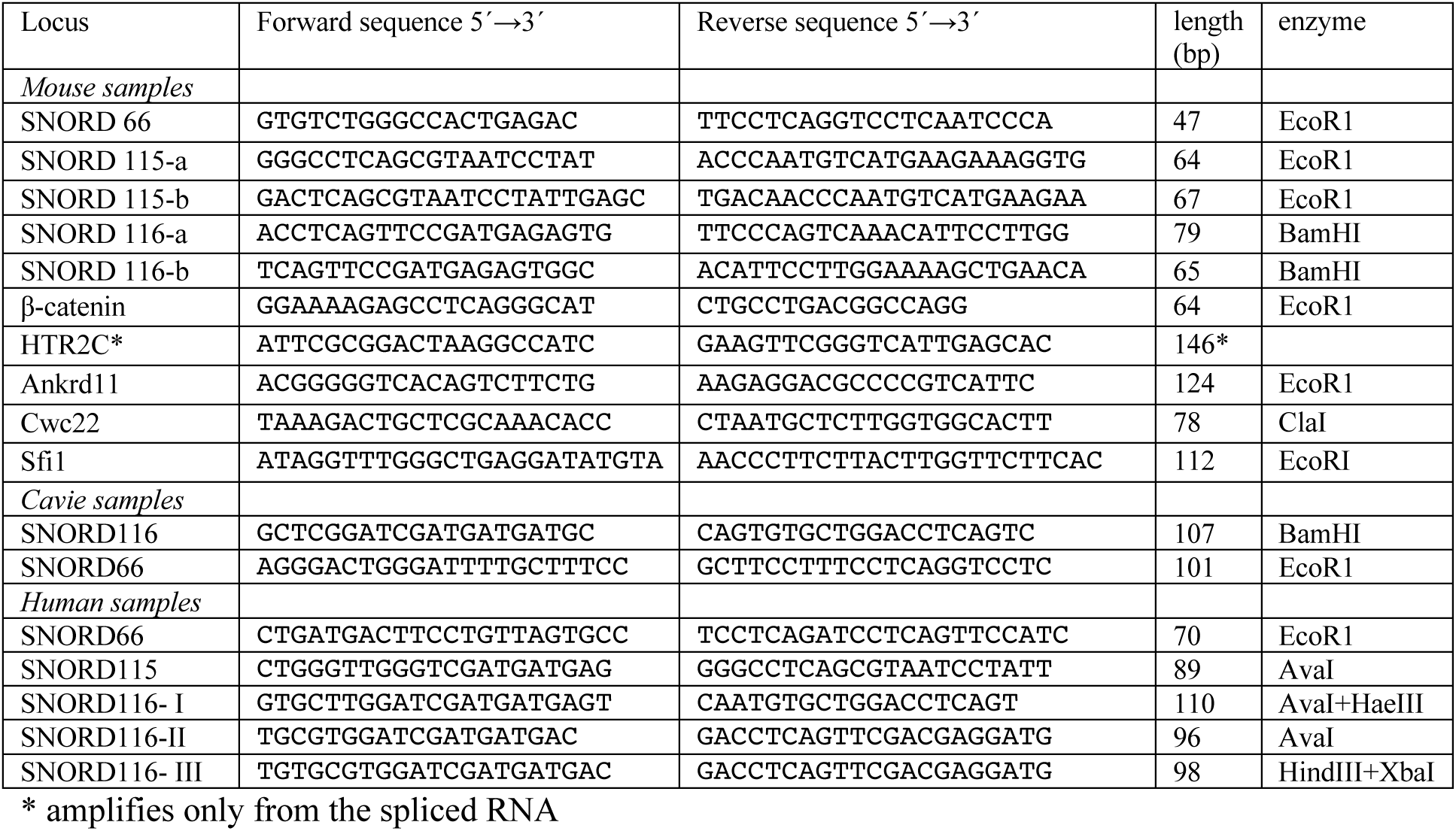
Primer sequences and enzymes for pre-digestion

## Discussion

The PWS region has been extensively studied in the past and it is clear that it is involved in a multitude of phenotypes. Given its implication in disease phenotypes in humans, a strong focus has been on loss of function studies, i.e. mostly deletions in the region, since these mimic the disease phenotypes. Our focus here was to look at the effects of natural variation caused by different copy numbers of two SNORD RNA clusters. These form two large tandemly repeated arrays that provide a technical challenge for their analysis. We show here that ddPCR is currently the only technique by which this can be solved. By comparing copy numbers with behavioral scores in a variety of populations and species, we were able to establish a link between these scores and copy number variation at the SNORD115 and SNORD116. This includes also an isogenic mouse strain, which is not expected to carry many other genetic variants (Zurita et al. 2011). Furthermore, RNA expression analysis reveals target genes that have previously been implicated in behavioral and other phenotypes known from PWS patients and knockout mice. Intriguingly, our data establish also a possible direct link to the skull shape phenotypes known in humans.

Further, our data show that new copy number alleles are generated every generation, which provides an explanation for the high individual variances observed between parents and offspring, as well as the low general heritability. At the same time, it can explain the observation that the behavioral scores tested for “personality” are repeatable within the lifetime of an individual. The new length variants are most likely generated in the germline, possibly only during meiosis, through unequal crossing over. Hence, a given individual carries a specific genetic variant, even though it would not necessarily inherit it to its offspring.

### Technical considerations

Our conclusions rely on the validity of the ddPCR data for measuring the repeat numbers. As discussed above, there are already a number of studies that show the reliability of ddPCR, including a validation by comparing ddPCR scores with sequencing depth data. In fact, if the ddPCR would be unreliable or too noisy, it would be very difficult to explain the observed correlations with SNORD copy numbers. Also, given that all copy number determination experiments were done blinded (i.e. the experimenter did not know the scores for the behavior tests), we can exclude observer bias.

Finally, a technical bias is highly unlikely, since SNORD115 and SNORD116 were amplified with different primer sets and since primer sets for two other high copy number CNV regions (*Sfi1* and *Cwc22*) did not yield any correlation.

There are also two further observations that make it unlikely that a technical error could cause the observed results. 1) We found an a priori unexpected correlation of SNORD115 and SNORD116 copy number in the ddPCR experiments. We have then re-checked previous data on genome read coverage for these regions (Pezer et al. 2015) and find the same correlation in the comparisons among the mouse populations. Hence, this can be considered as an additional validation of the ddPCR results. 2) The ddPCR results had suggested for the target gene expression data that the expression level of target exons correlated with copy number. Our subsequent RNASeq data allowed to test this prediction and we found it confirmed.

The choice of the behavioral assays could be another reason for concern. While these tests seem noisy, they constitute standard tests of behavioral biologists studying “personality”, i.e. they were found to be readily repeatable and consistent. We have paid extra attention that the tests were repeatable for given individuals over time. Forkosh et al. (Forkosh et al. 2019) have recently shown that the paradigm of screening for repeatable behavioral patterns among complex behaviors can be a proxy for “personality”. Our tests address one particular dimension of “personality”, namely anxiety. Interestingly, we find that the anxiety dimension of human personality tests via questionnaires shows also a correlation with SNORD copy numbers.

Finally, one can raise the concern that all our results are based on correlation only, rather than direct genetic interference. For example, we have not attempted to modify the number of active copies by a dedicated CRISPR/Cas mutagenesis approach. Rather, we looked at the effects of natural variation. Any attempt to do genetic manipulation would actually run into the problem of high natural variability between generations, i.e. it would not be possible to distinguish effects generated by CRISPR/Cas mutagenesis from the already existing normal variation, even in inbred strains. Hence, we opted to solve this verification problem by analyzing multiple different natural systems (i.e. animals form different populations and species). When one gets consistent results across these multiple different gnomic backgrounds, one can consider this as an independent confirmation that it is indeed the SNORD copy number variation that creates the effect rather than a consequence of something else in the genomic background of a given strain.

Finally, the PWS region has not shown an association in large scale GWA studies in mice that have included anxiety tests (Nicod et al. 2016; Parker et al. 2016). This would seem in contrast to the rather strong effects that we trace here. However, GWA studies rely on SNP typing and linkage to the typed SNPs. Such a linkage is not given when new length variants are generated at a high rate. Hence, the effects we trace here cannot be properly addressed via linkage analysis, but only via direct testing of copy number polymorphism, as we have done.

### Origin of variation

The high level of copy number variation in the SNORD115 and SNORD116 clusters can be ascribed to their tandem repeat organization, which should facilitate unequal crossover and thus length variation of the cluster. Both the family study in the wildtype mice, as well as the observation of many alleles in the otherwise isogenic strain C57BL/6J suggest an exceptional high rate of generation of new repeat number variants. C57BL/6J mice were originally generated from a single pair of mice in 1921 and have been kept as a well identifiable strain with very little SNP variation (Zurita et al. 2011). Still, we find at least 25 copy number variants for each SNORD cluster (when binned in distances of five) among the 57 animals that were provided by a single supplier (Jackson laboratories). While the exact number of generations that have passed since the founding of the strain is difficult to ascertain, it would not be more than a few hundred. Given that we find at least 25 alleles among them (and this can be considered to be a conservative estimate), this suggests at minimum the generation of one new variant every few generations. Imprinted regions are known to include hotspots of meiotic recombination (Paigen and Petkov 2010). However, such an extreme rate of change has not been described so far.

Another very unexpected finding is the covariation of repeat numbers between the SNORD115 and SNORD116 clusters. The repeat units of these clusters do not show any sequence similarity and they are separated by a non-repetitive region. We are not aware of any mechanism that could explain this covariation, but it is consistently found in all tested species. Given the overlapping regulation of the same behavioral pathways by the two clusters, it makes sense that they should show this covariation, but how this is achieved will require further studies.

### Target genes

Our transcript analysis data in response to SNORD115 copy number variation confirm the previously described interaction and regulation of the alternatively spliced exon Vb of the serotonin receptor *Htr2c* through this snoRNA (Kishore and Stamm 2006). We suggest that the corresponding predicted interaction of human SNORD116 with *Ankrd11* may also be a direct interaction, but we have so far not been able to clarify the mechanism. In contrast to the SNORD115 interaction, we do not find signs of RNA editing around the binding site in the transcriptome sequences. Instead, we find an influence on the expression level of the whole *Ankrd11* RNA. While this could be a consequence of alternative splicing, other regulatory mechanisms are not excluded. Note that C/D-box snoRNAS can function in many different ways (Falaleeva et al. 2017).

There are a number of mouse knockout lines that delete more or less large parts of the PWS region. However, only one knockout line exists that has removed specifically one SNORD gene cluster, namely SNORD116. The mice from this line show an early-onset postnatal growth deficiency, are deficient in motor learning and show slightly increased anxiety in one of the tests that were performed (Elevated Plus Maze) (Ding et al. 2008). The latter finding would be somewhat at odds with our results that the anxiety scores increase with the number of SNORD115 and SNORD116 copies.

However, we expect that the anxiety phenotypes are primarily regulated via the serotonin pathway that is affected by SNORD115 copy number and this was not controlled for in the (Ding et al. 2008) study. The SNORD116 copy number variation would be more relevant for the other associated phenotypes. In fact, a later analysis of a full deletion variant of this line has revealed further effects on feeding related pathways, as well as bone mineral density (Qi et al. 2016). These findings are in line with our findings of a disturbance of the multiple pathways that are regulated by *Ankrd11* as a direct target gene of SNORD116.

Interestingly, we find that not only behavior, but also craniofacial shape correlates with the SNORD copy number variation and the above reasoning would suggest that this is conveyed by SNORD116. For humans it has been suggested that there is indeed a link between personality and facial characteristics (Kramer and Ward 2010), but a possible causality of these observations was left open. Our data suggest such causality for mice. But given that mice are essentially nocturnal animals that communicate mostly via scents and ultrasonic vocalization (von Merten et al. 2014), it is unclear whether they would even recognize different craniofacial shapes among their conspecifics. However, the cranial shape changes could reflect a general osteogenic effect on the whole bone system that could be of relevance for being combined with the behavioral tendency. For example, bold animals might profit from stronger bone structures, in case they get more involved in fights. However, this connection will need further study, especially since the SNORD116 expression is confined to the brain. Qi et al. (Qi et al. 2016) have suggested that a general osteogenic effect may be mediated via the metabolic pathways that are regulated by *Ankrd11*.

### Implications for the genetics of anxiety traits

Our findings can resolve a long-standing controversy about the genetics and plasticity of anxiety or “personality” traits. Our data suggest that these traits are indeed genetically specified, but through a hypervariable locus that would escape conventional genetic mapping approaches. Hence, it is not necessary to invoke developmental or secondary epigenetic effects in explaining variance in such traits, given that there is a mechanistic pathway of modulation of target RNA expression through different concentrations of snoRNAs. However, such alternative mechanisms may have additional contributions. In future studies on such traits it would seem advisable to treat SNORD copy number variation as a covariate to assess whether other mechanisms exist as well.

The fact that we find significant regressions between behavioral scores and SNORD copy numbers also in other rodents and even in humans, implies that the system is conserved throughout mammals. In humans, we observed this within a psychiatrically healthy control group, where we find that extremes of personality traits for anxiety and activity are associated with significant differences in copy numbers. The effects are somewhat weaker than for the rodents, but humans are evidently also more subjected to environmental influences than the animals that were bred under controlled conditions. Most intriguingly, however, the effect in humans is in the opposite direction, i.e. more anxious individuals harbor lower copy numbers. This would suggest that the copy number variation acts only as a general regulator, but the actual behavioral consequences are modified by downstream pathways.

### Evolutionary implications

One can raise the question of how and why such a hypervariable system could have evolved and how it is maintained. This question has in fact been posed since a long time and there have been a number of attempts to propose evolutionary models that could explain the large variability in behavioral traits. These include drift effects in a neutral context (Tooby and Cosmides 1990), mutation-selection balance (Zhang and Hill 2005) or balancing selection processes (Dingemanse and Wolf 2010; Penke and Jokela 2016). But none of these models has taken the possibility into account that a hypervariable locus could control this variation. Hence, our results establish a new paradigm in the regulation of behavioral variance, namely the involvement of dosage changes of a regulatory RNA through recurring copy number mutations.

## Acknowledgements

We thank Chen Xie for his valuable advice with transcriptome analysis, Stefan Dennenmoser, Cemalettin Bekpen and Guy Reeves for their advice on SNORDs copy number analysis, Sven Künzel for sequencing, Ellen McConnell, Nicole Thomsen and Cornelia Burghardt for their great lab-related support, Elke Blohm-Sievers for her help in CT scanning, the entire mouse team at MPI for evolutionary biology, in particular Christine Pfeifle, Susanne Holz and Camilo Medina for taking care of the mice in this study and Heike Harre and Fikri Tuğberk Kara for their help with behavioral tests and brain dissection. Animals were kept according to FELASA (Federation of European Laboratory Animal Science Association) guidelines, with the permit from the Veterinäramt Kreis Plön: 1401-144/PLÖ-004697and the Veterinäramt Bielefeld for cavies, respectively. The experiments involving the behavioral analyses was approved and registered under V244-71173/2015 through the local authorities. The animal welfare officer was informed about the sacrifice of the animals for the molecular analyses. The study on humans was approved by the ethics committee of the Ludwig-Maximilians-University (LMU) in Munich and written informed consent was obtained from all subjects.

## Methods

### Mouse strains

Wildtype mice (*Mus musculus domesticus* and *Apodemus uralensis*) used in this study were offspring of mice that originated from wild populations sampled in the Massif Central region of France (MC), the Cologne/Bonn region of Germany (CB) and Kazhakstan, Almaty (*Apodemus*) and then held under outbreeding conditions at the Max-Planck-Institute for Evolutionary Biology in Plön. The C57BL/6J inbred strain (30 male and 30 female) was purchased at the age of 3 weeks from Charles River Laboratories, Research Models and Services, Germany GmbH (Sandhofer Weg 7, 97633 Sulzfeld). These inbred mice were not siblings.

All mice were bred in type III cages (Bioscape, Germany) and were weaned at the age of 3 weeks. At weaning, the sexes were separated. Males were housed together with brothers or in individual cages. Females were housed in sister groups to a maximum of 5 mice per cage. Enrichment, including wood wool, toilet paper, egg cartons and a spinning wheel (Plexx, Netherland), was provided in each cage. Mice were fed standard diet 1324 (Altromin, Germany) and provided water ad libitum. Housing was at 20–24°C, 50–65% humidity and on a 12:12 light-dark schedule with lights on at 7 am. While the C57BL/6J and the *Apodemus* mice and MC mice used in family study were kept under identical conditions throughout the experiment, the CB and MC mice in the first part of study were released into semi-natural conditions at an age of 3-4 months. Details of this experiment are provided in the PhD thesis of Rebecca Krebs which is available at https://macau.uni-kiel.de/receive/dissertation_diss_00022305.

### Wild cavies

Details on the behavioral work with wild cavies are described in previous papers (Guenther et al. 2014; Guenther and Trillmich 2015). Wild cavies were descendants from a population of wild animals caught in Uruguay in 2003 and 2007. Experimental animals were bred in outdoor enclosures consisting of a roofed hut (2m^2^) and an outdoor compartment (14m^2^) in the Department of Animal Behaviour at Bielefeld University, Germany. They experienced natural temperature and photoperiodic conditions. Food (hay and guinea pig chow, Höveler, Germany) and water was provided ad libitum and supplemented with fresh green (e.g. carrots, bellpepper or apples) three times a week.

At weaning (21 days of age), animals were individually marked and transferred into indoor standard housing conditions. Indoors, animals were kept in unrelated same-sex groups of four individuals each and housed in 1.6m^2^ enclosures. Each enclosure was equipped with three shelters, three feeding dispensers and three water bottles. Animals were kept under natural light conditions with additional artificial light from 6 am to 7 pm, at constant temperatures of 20 ± 2 °C. The feeding regime was the same as under outdoor conditions. Animals used for this study were all unrelated. In total, 10 males and 9 females were used for the experiments. Since wild cavies do not show sex-differences in the expression of anxiety-related behaviors (Guenther and Trillmich 2015), we did not include “sex” as a factor in statistical analyses.

### Human cohort and evaluation of personality traits

541 healthy controls were randomly selected from the Munich registry of residents and interviewed for the presence of DSM-IV anxiety, affective, somatoform, eating, alcohol dependence, drug abuse or dependence, disorders using a modified version of the Munich Composite International Diagnostic Interview (Wittchen and Pfister 1997) at the MPIP. Only individuals negative for the above-named disorders were included in the context of a large study on the genetics of major depression using EDTA blood as the source for DNA (see (Heck et al. 2009) for more detail).

### DNA and RNA extraction and cDNA synthesis

All dissections were done following standardized protocols and personal instructions. Prepared tissues were immediately frozen and kept at −70 ° until DNA/RNA preparation.

DNA extraction was performed according to a standard salt extraction protocol. Briefly, samples were lysed by using HOM buffer (80 mM EDTA, 100 mM Tris and 0.5 % SDS) with Proteinase K (0.2 mg/mL) for 16 hours in Thermomixer (Eppendorf, Germany) at 55°C. 500 μL sodium chloride (4.5 M) was added to each sample and was incubated on ice for 10 minutes. Then chloroform was added, mixed and spun for 10 minutes at 10,000 rpm. The upper aqueous phase was separated, mixed well with Isopropanol (0.7 volume) and spun for 10 minutes at 13,000 rpm. The pellet was washed with Ethanol (70 %), air dried and dissolved in TE-buffer (10 mM Tris, 0.1 mM EDTA). DNA concentration was measured on the Nano Drop 3300 Fluorospectrometer using Quant-iT dsDNA BR Assay kit (Invitrogen) reagent.

RNA extraction (from whole brain) was done by using Trizol reagent. 1mL Trizol was added to each sample. Then the samples were lysed by Tissue lyser II (QIAGEN, Germany) at 30 Hertz for 5 minutes. Homogenized samples were incubated at room temperature for 5 minutes. 200μL chloroform (per 1 mL TRIzol) was added to each sample, shook vigorously by hand 15 seconds, followed by 3 minutes incubation at room temperature and spun at 12,000g for 15 minutes at 4°C. The aqueous phase was transferred to a new tube and 0.5 volumes Isopropanol was added, incubated at room temperature for 10 minutes and spun at 12,000g at 4°C. The supernatant was removed and the pellet was washed with 75% EtOH (made with RNAse-free water). Samples were mixed by hand several times and then spun at 7,500g for 5 minutes at 4°C. The supernatant was removed and the pellet dried shortly at room temperature, dissolved in 200μl RNAse free water and stored at −20°C for overnight. An equal volume of LiCL (5M) was added to the crude RNA extract, mixed by hand and incubated for one hour at −20°C. Samples were spun at 16,000g for 30 minutes. The supernatant was removed; samples were washed twice with EtOH 70% and spun at 10,000 at 4°C. The pellet was dried at room temperature, dissolved in RNAse free water and kept for long-term storage at −70°C.

The quality of the RNA samples was measured with Bio-Analyzer chips and samples with RIN values below 7.5 were discarded. cDNA was synthesized using the MMLV High Performance Reverse Transcriptase kit according to the instructions of the supplier (epicenter, an Illumina company).

### Small RNA extraction and cDNA synthesis

Total RNA was extracted from the whole brain by using the mirVana miRNA Isolation Kit, which enriches small RNAs. The quality of the RNA was measured with BioAnalyzer chips and samples with RIN values below 8 were discarded. Illumina® TruSeq® Small RNA Library Prep kit was used for small RNA cDNA synthesis. The protocol in the Illumina kit takes advantage of the common natural structure in most known small RNA molecules. Most mature small RNAs have a 5’-phosphate and a 3’-hydroxyl group. So, the Illumina adapters in this kit are directly ligated to these small RNAs. Then Reverse Transcriptase with a primer for this adaptor was used to synthesize cDNA from small RNA. 1 μg of total RNA which was enriched in small RNA was used as input. To control for DNA contamination, we included a tube with all ingredients apart of the reverse transcriptase. The results obtained from this tube were used as gDNA contamination to normalize the gene expression data for each sample. The data provided in Suppl. file 12 show that gDNA contamination is expected to be very small for each sample.

### RNAseq analysis

Poly-A^+^ RNA was used for cDNA synthesis and Illumina library preparation by using the Truseq stranded RNA HT kit. The libraries passing quality control were subjected to sequencing on an Illumina NextSeq 500 sequencing system and paired-end reads were 150 bases long. Raw sequence reads were quality trimmed using Trimmomatic (Bolger et al. 2014). The quality trimming was performed base wise, removing bases below quality score of 20 (Q20), and keeping reads whose average quality was of at least Q60. Reads were mapped to the mouse mm10 reference genome by using Hisat2 (Kim et al. 2015). Between 20.1 to 28.7 million mapped reads were obtained per sample (average 23.8 million). Differential expression analysis was performed with the DESeq2 package (Love et al. 2014) in the R environment. To perform the alternatively spliced isoforms analysis, BAM files from Hisat2 were used as input for SAMtools (Li et al. 2009) by using option –c for total read from each exon and option –q 60 for total read of each sample.

### Droplet digital PCR

Digital PCR is a method enabling absolute quantification of DNA targets without the need to construct a calibration curve as used in qPCR (Zhao et al. 2016). It requires a reference to calculate gene copy number and to normalize gene expression level. For the latter we used β-catenin for mRNAs and SNORD66 for standardizing SNORD RNAs. SNORD 66 is a single copy gene located in an intron of the eukaryotic translation initiation factor 4 and was therefore used also as reference gene for copy number calculations. All reactions were done as technical triplicates which yielded highly congruent results (S1 Table). Averages of the triplicates were used to calculate copy numbers. Values for copy number represent the total number of copies per diploid genome. The values for expression represent the number of RNA copies per µL of input material.

Primers (Table 3) were designed aiming for a balanced GC content <50% and with low potential for primer dimer structure. All primers were verified that they amplify only the expected length fragment.

The SNORD115/116 primers were placed within the coding sequences of the respective snoRNAs. Two pairs were tested for each for an initial set of 19 animals and found to yield highly congruent results (R^2^=0.99, S3 Table). Hence, we used only the first primer set (labelled with -a) for the rest of the analyses. To test for biological replicability, we obtained ear clip samples from 41 four-week old animals and later (at 14 weeks) brain samples from the same animals. Copy number determination was highly congruent between these samples (R^2^=0.99, S3 Table).

The QX100™ Droplet Digital™ PCR System (Bio-Rad, Hercules, CA, USA) was used in this study according to the manufacturer’s instructions. Briefly, fluorescent PCR reactions for each sample were prepared in a 23 μL volume containing 12μL 2 X EvaGreen supermixes, 200nM of each forward and reverse primers, DNA or cDNA (1ng in mice study,10ng in Guinea pig and 5ng in human study) and water. For tandem copy separation, sample viscosity reduction and improved template accessibility, DNA digestion was done by different restriction enzymes (Table 3, Thermo Fisher Scientific, Germany). 5 units of restriction enzymes were added to each sample. The samples were kept 20 minutes at room temperature for complete digestion.

Droplets were generated using a Droplet Generator (DG) with an 8-channel DG8 cartridge and cartridge holder with 70 μL of DG oil/well, 20 μL of fluorescent PCR reaction mixture and a DG8 gasket. The prepared droplets were transferred to corresponding wells of a 96-well PCR plate (Eppendorf, Germany). The PCR plate was subsequently heat-sealed with pierceable foil using a PX1™ PCR plate sealer (Bio-Rad) and then amplified in a LifeEco thermal cycler (Bioer, China). The thermocycling protocol was: initial denaturation at 95°C for 5 min, then 40 cycles of denaturation at 95°C for 30 s, annealing at 60°C for 45 s and, finally, incubation first at 4°C for 5 min and then at 90°C for 5 min. After cycling, the 96-well plate was fixed into a plate holder and placed into the Droplet Reader. Droplets of each sample were analyzed sequentially and fluorescent signals of each droplet were measured individually by a detector. Copy numbers were calculated according to the procedures suggested by BioRad using QuantaSoft 1.7.

## Behavioral Tests

### Mouse strains

Behavioral tests were performed on the *M. m. domesticus* mice starting at an age of 24 weeks, on theC57BL/6J inbred strain at the age of 16 weeks and on the *A. uralensis* mice at the age of 18 weeks. The testing included an Elevated Plus Maze and an Open Field, all adapted towards the use of wild mice to avoid escape. Test setups were cleaned with 30% ethanol between individual tests. All tests were done during the day (between 8 am to 1pm) with constant artificial light of about 300 lux.

The Elevated Plus Maze consisted of four arms 1m above ground, each 50 cm long and a 10×10 cm neutral area in the middle. Two of the arms were made of clear Plexiglas, indicating the unsafe zone and two were made of grey PVC, indicating a safe zone. The floor was made of white PVC. Mice were placed in the center and the behavior of each mouse was monitored for 5 min. During this experiment, the time spent in the dark and light arms were measured, as well as the speed and distance travelled.

For the Open Field test, mice were placed in a 60×60 cm arena with 60 cm high white PVC walls and they were allowed to explore it for 5 min. The speed of the mouse, the distance travelled and time spent within 10 centimeters off the wall vs in the central area were measured.

The behavioral tests were filmed using a TSE camera (TSE system, Germany). To score the videos from each test, all the videos were transferred to Videomot2 system (TSE system, Germany). Mice were detected by the software in 3 points (head/center/tail base tracking) and then the software automatically generates the numerical data for the above-mentioned measurements.

### Wild cavies

The behavioral tests on the wild cavies (*Cavia aperea*) were conducted as described in (Guenther and Trillmich 2015). The tests started when individuals were 55 ± 3 days old. In short, to measure struggle docility, the animal was turned on its back and held in the hand of an observer. For 30 s, the time the animal struggled to actively escape that situation was recorded. To measure open field anxiety, an Open Field test was used. This consisted of a 1m^2^ arena and behaviour was scored based on the videos from a camera on top of the field. Light conditions were artificial and constant around 250 lux.

Experiments were conducted between 9-12 am or 3-5 pm, as this was shown to have no systematic influence on the behaviour (Guenther et al. 2014). The distance moved (cm) when individuals were exposed to an open field for 20 min, was scored. The first 10 min, a semi-transparent shelter (20cm x20cm with four exits) was present in the arena under which animals could hide. For the second 10 min, this shelter was lifted out of the arena. Individuals which are anxious in this situation freeze and move about only little while individuals which are not anxious run around and explore the situation (Guenther and Trillmich 2015). Hence, for wild cavies, high values equal low anxiety.

### Humans

Five hundred and forty-one individuals also completed the Tridimensional Personality Questionnaire (TPQ), a validated personality measure (Cloninger et al. 1991; Weyers et al. 1995). To match the behavioral assessments in mice, single items of the TPQ reflecting anxiety and activity were used to select individuals on the extremes of this distribution. Individuals can score from 0 to 6 on the activity and anxiety scale, respectively and their sum was used to identify the extreme groups, while matching for age and sex. The high anxiety and activity group (N = 48) had a mean additive score of 7.35 (SD 1.04). The low anxiety and activity group (N = 48) had a mean additive score of 0.73 (SD 0.49). The mean age of the high anxiety group was 42.3 year and 41.6 years for the low anxiety group with 33.3% and 62.5% females, respectively.

## Statistical analyses

### Repeatability of behavioral measures

As pointed out in the introduction, one of the cornerstones when aiming to describe *consistent* individual differences in behavior, is the validation of the temporal consistency of the behavioral trait. Commonly, an index called “repeatability” is used to describe the temporal consistency. Repeatability reflects the amount of variation between individuals compared to the amount of all phenotypic variation in the population (including also within-individual variation and measurement error).

To estimate temporal consistency for our animals, we conducted repeatability analyses for each of our three mouse model systems separately (i.e.: C57BL/6J, the wild mice MC population used in the family study and *Apodemus*). Each model included three measurements per behavioral trait in total, with 4 weeks interval between them. The CB and MC mice in the first part of study were part of an experiment in semi-natural environments. They were tested 8 weeks apart of each other.

Statistical analyses were carried out using R 3.3.3 and R 3.3.2. *Cavia aperea* and the CB and MC mice in the first part of study were a subset of mice that have been used in other projects and had already been analyzed for repeatability, so we did not include them for repeatability assessment in this study. For C57BL/6J, the wild mice MC population used in the family study mice and *Apodemus*, repeatability was calculated using the “rptR” package (Stoffel et al. 2017). Normally distributed data were calculated using rpt (ANOVA based) and for count data rpt.poisGLMM was used.

### Linear Regression analyses SNORD copy number variation and behavioral variables

To create a regression model for SNORD copy number and behavioral variables, the lm function in R. 3.3.3 and R 3.3.2 was used and population origin, sex and age were modelled as main factors. The average from triplicate measurements in ddPCR was used as SNORD copy number for each individual and the direct scores from first time measurement of each behavioral test were used as behavioral score in this study. p-values ≤ 0.05 were considered to be significant.

### Mixed-model regression SNORD copy number variation and behavioral variables

Univariate mixed models for each repeatable behavioral measurement in the behavioral experiments (for Open Field: time at the wall; for Elevated Plus Maze: time in the dark arm) were constructed using lmer function in R. 3.3.3 and R 3.3.2, including animal ID as random effect and triplicate measurements of SNORDs copy number from ddPCR as fixed effects. This analysis has been performed to account for the potential differences over the course of the behavioral experiments (three times behavioral measurement) as well as technical error from triplicate measurements from ddPCR for SNORDs copy number calculation. R^2^ was calculated to assess the proportion of variance explained by the model using r.squaredGLMM function from MuMIn package (Barton K., 2019 - https://CRAN.R-project.org/package=MuMIn). The P-value was calculated using sjstat package (Lüdecke 2019). p-values ≤ 0.05 were considered to be significant.

### Craniofacial shape analysis

Landmarks for the morphometric analysis were obtained as described previously (Pallares et al. 2015). Briefly, mouse heads were scanned using a computer tomograph (micro-CT—vivaCT 40; Scanco, Bruettisellen, Switzerland) at a resolution of 48 cross-sections per millimeter. 44 three-dimensional landmarks were positioned in the skull (the landmarks are described in (Pallares et al. 2015)). The raw 3D landmark coordinates obtained in TINA tool were exported to MorphoJ (Klingenberg 2011) for further morphometric analyses. The symmetric component of the skull was obtained through a mirror image of the landmark configuration of each individual was generated, and a full GPA was performed with the original and mirror configurations. Again, the resulting configurations were averaged to obtain the symmetric component of shape variation. The new landmark coordinates generated by the GPA are called “Procrustes coordinates”. The multivariate regression of the shape data (Procrustes coordinates) on SNORD116 copy number was carried out via function procD.lm from the geomorph R package v3.0.7 (Adams and Otarola-Castillo 2013).

To visualize the patterns of predicted shape differences for minimum and maximum values of SNORD116 copy number, the closest specimen to the overall mean shape in multidimensional space was first identified via the function ‘findMeanSpec’ in the geomorph package v3.0.7 (Adams and Otarola-Castillo 2013) and a 3D CT-scan of the specimen closest to the mean shape was warped to this overall mean shape using the Thin Plate Spline (TPS) method via the function ‘tps3d’ in R-package Morpho v2.6 and using warping and heatmap functions from the R-package Rvcg v0.18 (Schlager 2017) as well as the color palette from R-package viridis v0.5.1 (https://CRAN.R-project.org/package=viridis).

## Supplementary Figures

**S1 Figure:**
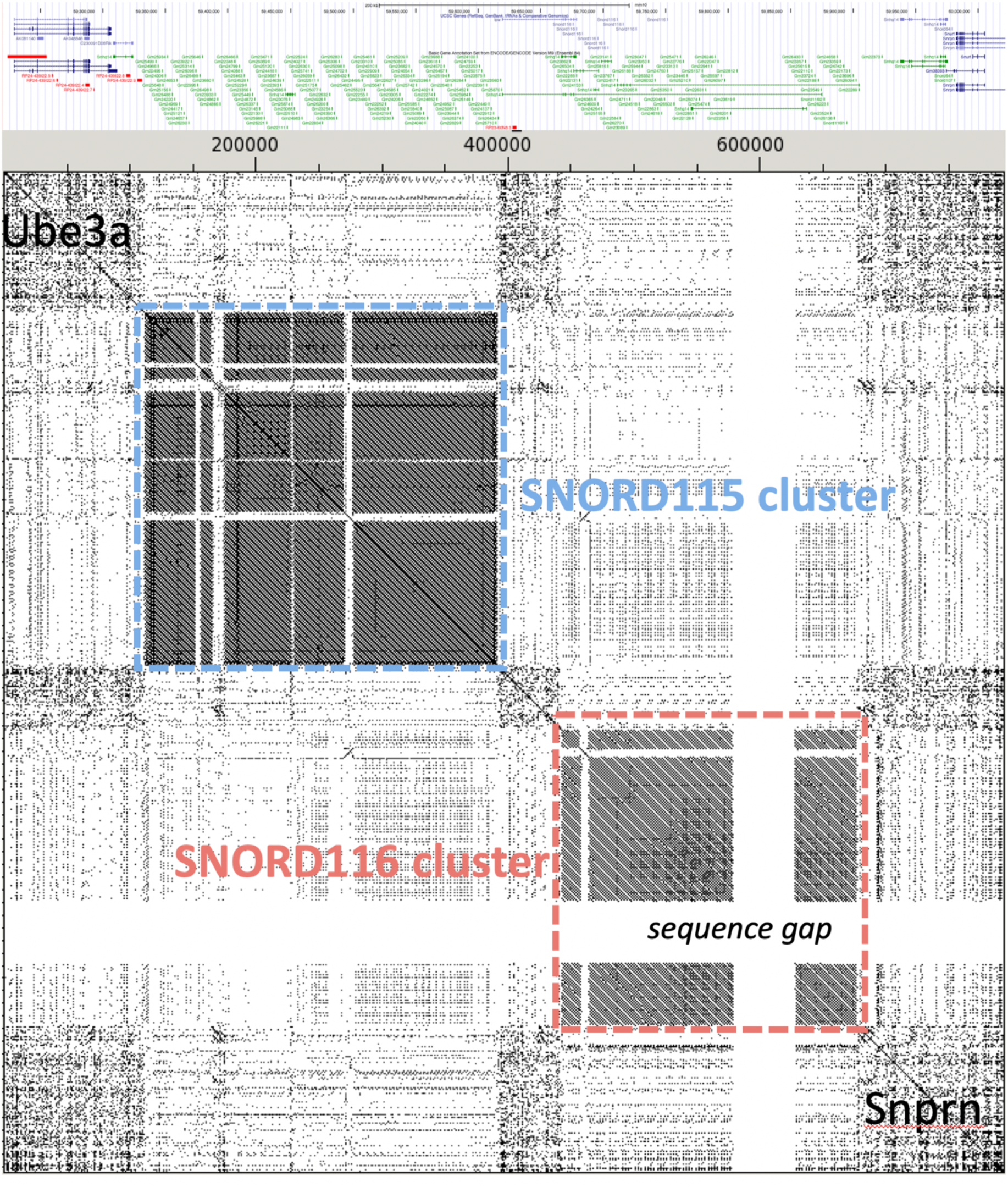
SNORD115 and SNORD116 tandem repeat clusters. Dotplot representation of the PWS region between *Ube3a* and *Snrpn* matched against itself. Sequence regions with matches appear as diagonals, tandem repeats as clusters of diagonals. The corresponding annotations from the UCSC browser are shown at the top. The Snord115 and SNORD116 clusters are boxed, the sequence gap (i.e. a 50kb region filled with N in the reference sequence) is indicated.

**S2 Figure:**
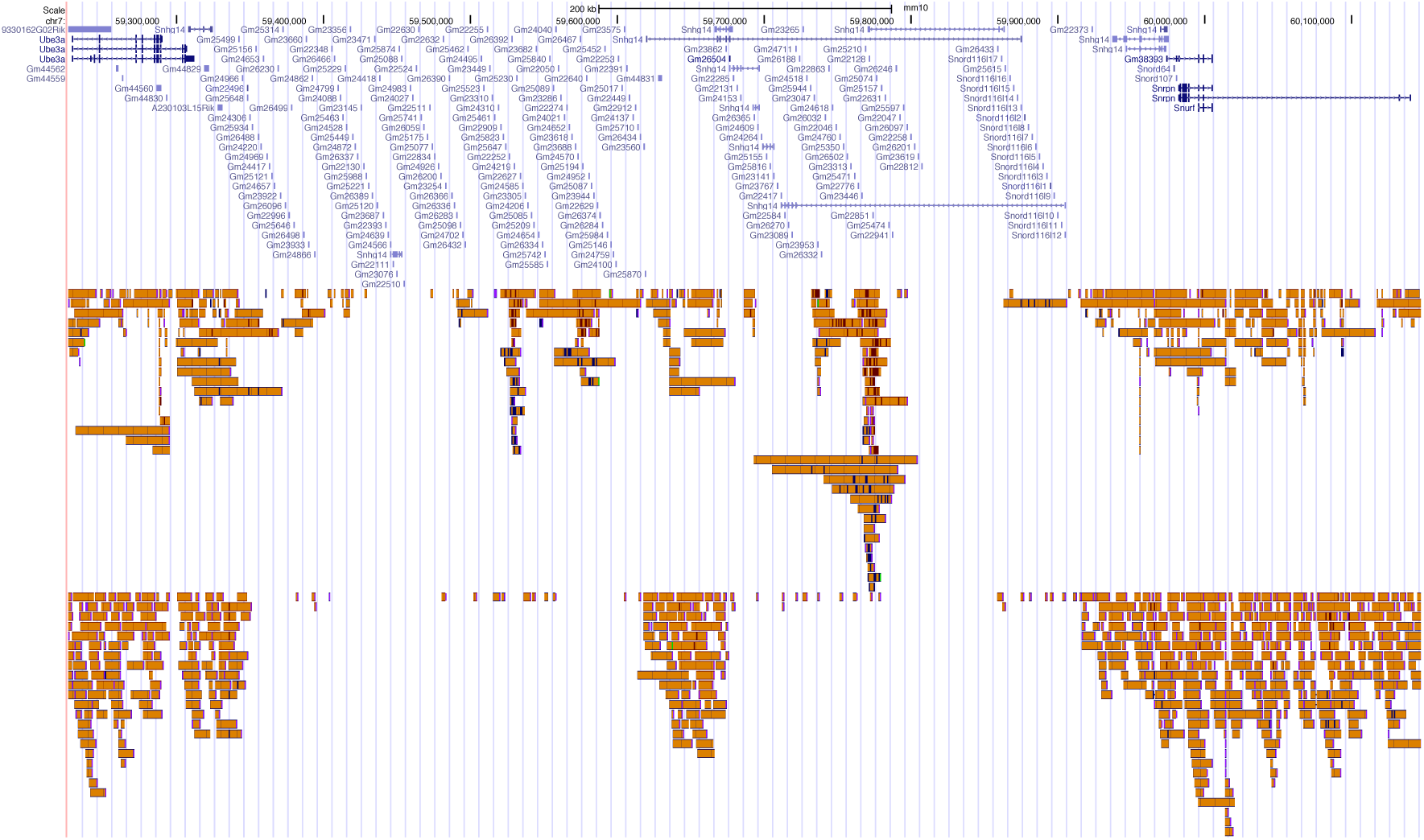
Long read coverage comparisons. UCSC browser tracks showing the region between *Ube3a* and *Snrpn* with the corresponding reads obtained from Oxford Nanopore (top track) and PacBio (bottom track) sequencing of whole genomes (compiled from several runs). Each read is represented by an orange bar showing the extent of its coverage. Note that the mapping algorithm does not place reads in regions of internal repeats unless there are specific additional SNPs that allow unequivocal placements. This explains the relative paucity of reads in the SNORD115 and SNORD116 tandem repeat clusters.

**S3 Figure.**
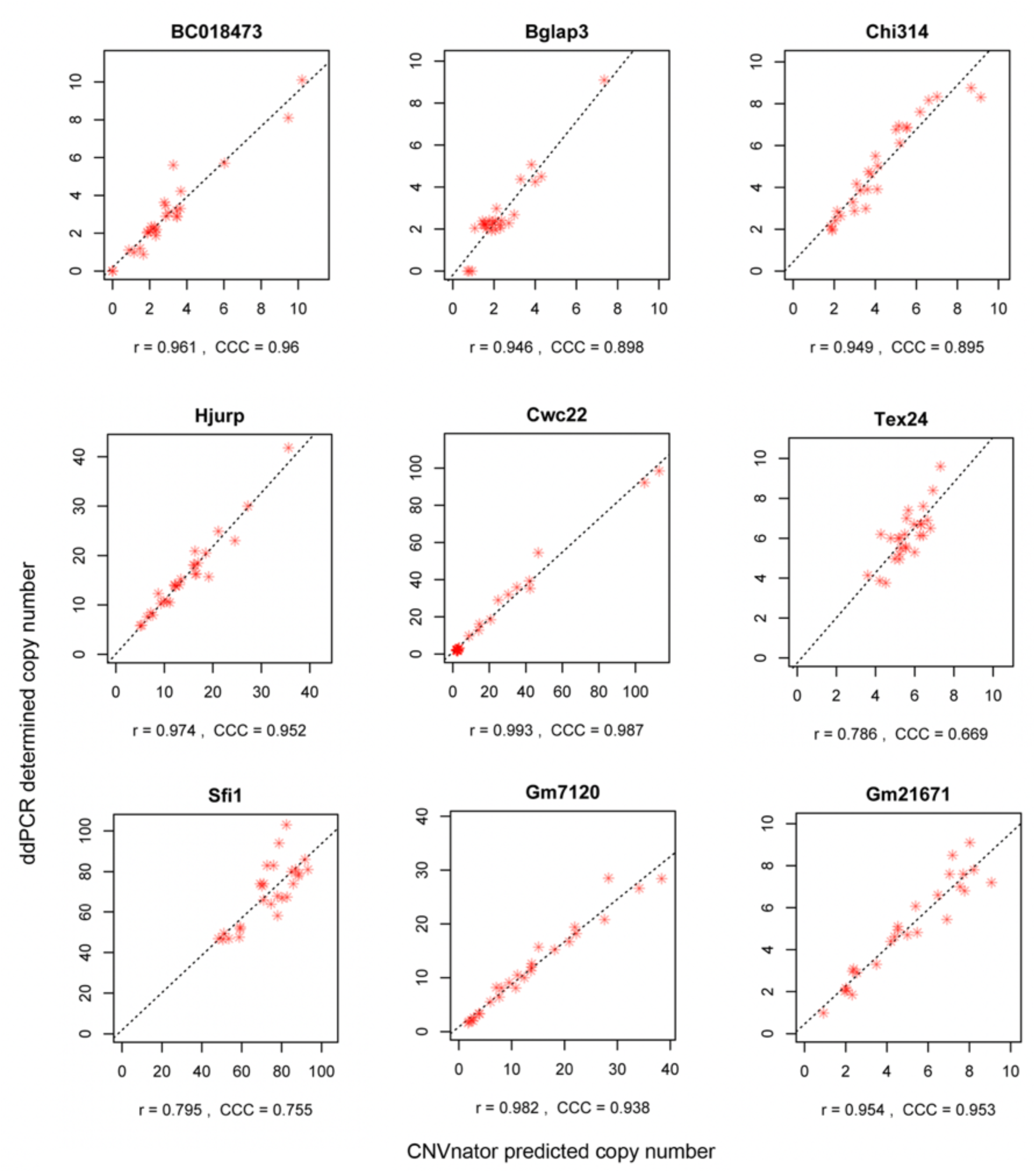
Comparison of copy numbers at tandem repeat loci determined by sequencing read depth versus digital PCR. Assays for 9 loci with copy numbers of 10 or higher were designed and used to screen 27 individuals (red asterisks) for which read depth information was available from whole genome sequencing; data taken from Pezer et al. (Genome Res 25, 1114-1124, 2015). The results of ddPCR runs (y axis) are plotted against copy numbers predicted with CNVnator based on sequencing read depth (x axis) for the amplicon regions. Fitted regression lines are shown dashed; *r* is Pearson’s correlation coefficient; CCC is Lin’s Concordance Correlation Coefficient.

**S4 Figure.**
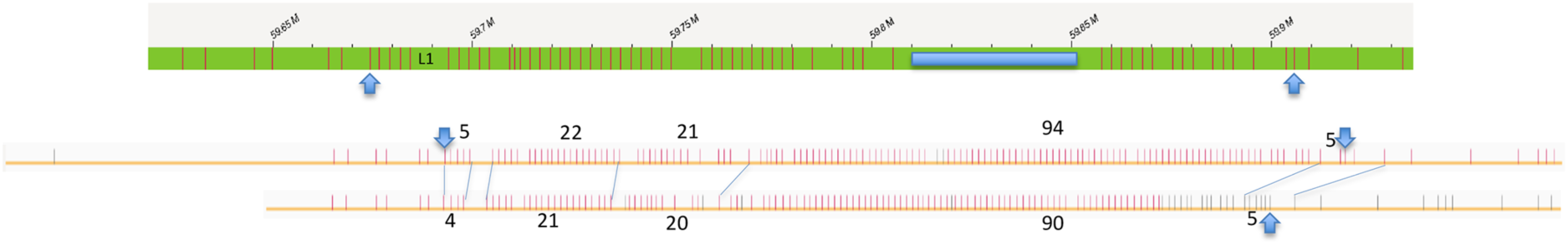
Optical mapping analysis of the SNORD116 region. Optical mapping was done following the standard procedures on a Bionano Saphyr instrument using Nt.BspQI, which labels GCTCTTC strings in the DNA. This string occurs once in the SNORD116 repeat consensus sequence. Fragments that span the whole region of the SNORD116 cluster plus the adjacent regions were manually retrieved from the fragment display function of the instrument software. The green bar at the top represents the predicted labelling positions for the restriction enzyme from the mouse reference genome. The tandem repeat region (marked by the arrow heads) becomes apparent by the regular spacing of sites, which are, however, not perfect. The string includes at one end an inserted L1 element that interrupts this spacing and which is marked in the bar for the reference sequence. We find this same interruption in the recorded fragments and this serves as a suitable marker to align the fragments with the reference sequence. There are further interruption of the regular pattern that are due to repeat copies that lack the Nt.BspQI site. The reference sequence includes also a 50kb gap filled with N (marked in blue). The lines below the reference bar represent two fragments that are interpreted to constitute the two alleles of this individual. Visible gaps are used to align them (blue line connectors) and count the number of repeats (annotated numbers). Note that the repeat counts consider the gaps in the regular string as variants that have no Nt.BspQI site or labelling drop outs, i.e. they are added to the total numbers. Allele 1 of this mouse shows 147 copies, allele 2 shows 140 copies. The ddPCR measurement for this mouse was 293 copies, which is well within the range of the total of 287 predicted copies from the optical mapping.

**S5 Figure.**
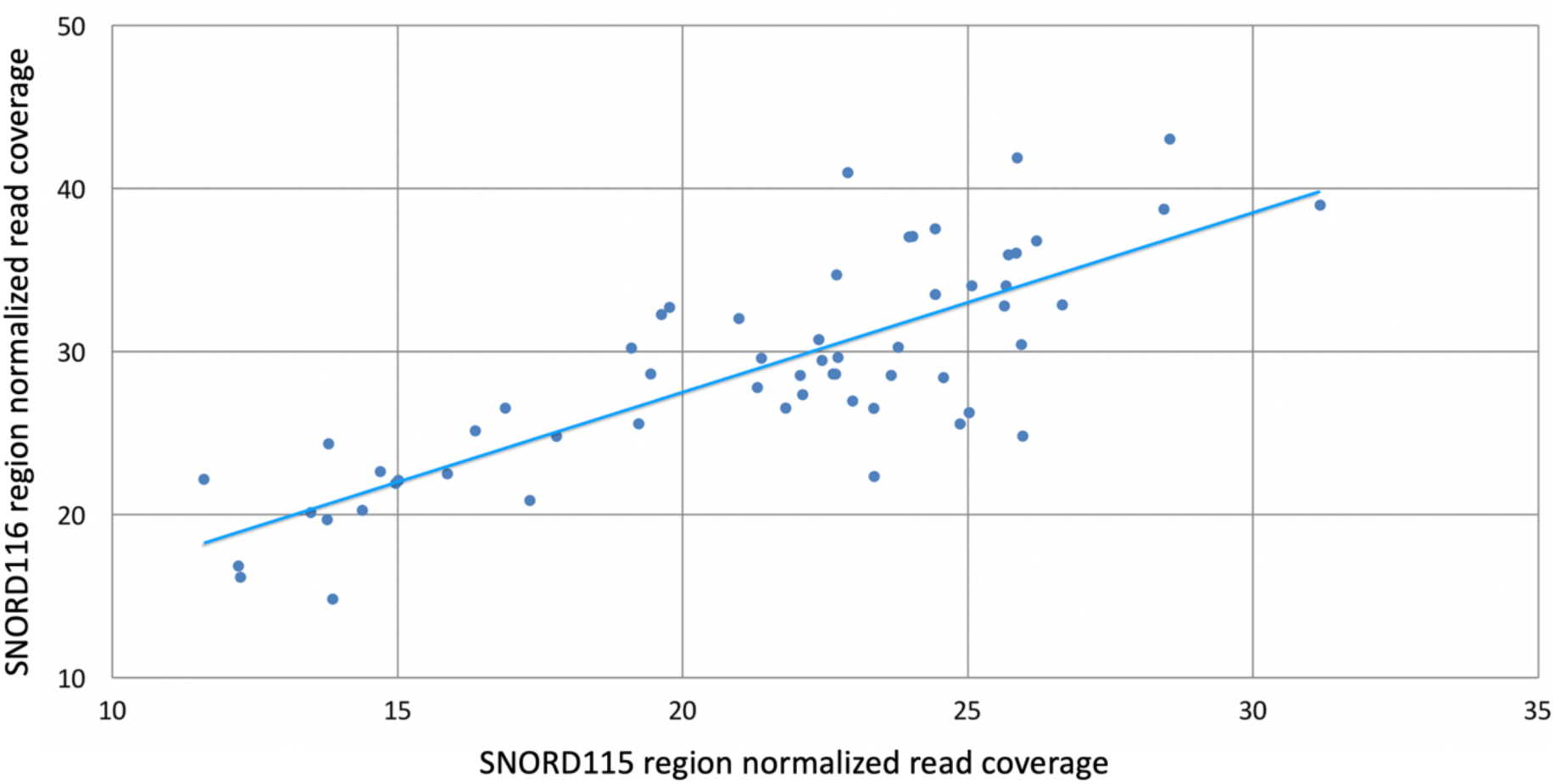
Correlation of read coverage between the SNORD115 and SNORD116 regions based on whole genome sequencing data. Genome sequencing reads from individuals of Mus musculus populations, including the three subspecies (Harr et al. 2016) were mapped against the house mouse reference genome mm10 with ’ngm’ (Sedlazeck et al. 2013), followed by sorting, marking and removing duplicates with the picard software suite (https://broadinstitute.github.io/picard/). The resulting bam files can be obtained via http://www.user.gwdg.de/~evolbio/evolgen/wildmouse/introgression/ngm/bam/. We used the ’genomeCoverageBed’ tool from the bedtools software suite v2.26.0 (Quinlan and Hall 2010) with the option ’bga’ to obtain site-specific coverage across the snord115 (chr7:59,340,775-59,620,255) and snord116 (chr7:59,671,705-59,905,815) region. Subsequently, the coverage files were used to calculate the mean coverage per site in the respective genome regions. These mean coverages were then normalized according to the total reads obtained from each genome.

**S6 Figure.**
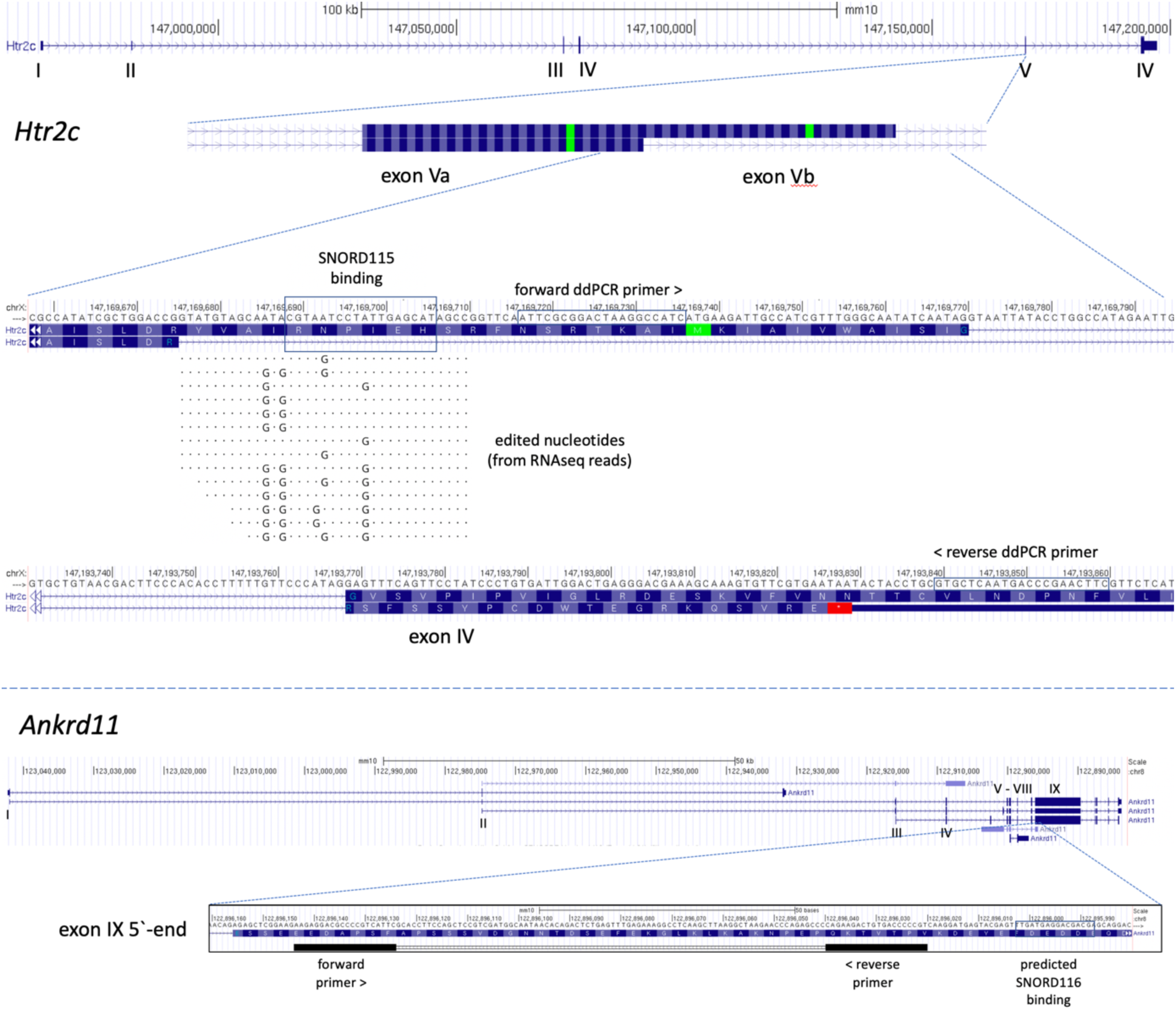
Structures of the *Htr2C* and Ankrd11 genes. The structure details with exon arrangements are taken from the UCSC browser. Exon numbers are indicated in roman numerals. For *Htr2c*, exon V and VI are enlarged, for *Ankrd11* exon IX is enlarged. SNORD binding sites and primer sites for the ddPCR experiment for determining copy numbers are marked with boxes. Further, we added a read alignment from RNASeq reads to show the edited sites in *Htr2c*.

## Supplementary Tables

**S1 Table.**

|  | A | B | C | D | E | F | G | H | I | J | K | L | M |
| --- | --- | --- | --- | --- | --- | --- | --- | --- | --- | --- | --- | --- | --- |
| 1 | SNORD115 and SNORD116 copy numbers for *Mus* strains and stocks |  |  |  |  |  |  |  |  |  |  |  |  |
| 2 |  |  |  |  |  |  |  |  |  |  |  |  |  |
| 3 |  |  |  | SNORD115 copy numbers |  |  |  |  |  | SNORD116 copy numbers |  |  |  |
| 4 | MouseID | Populations / strain | sex | SNORD115<br>(replicate1) | SNORD115<br>(replicate2) | SNORD115<br>(replicate3) | average | SD | SNORD116<br>(replicate1) | SNORD116<br>(replicate2) | SNORD116<br>(replicate3) | average | SD |
| 77 | 50046761 | Mouse inbred strain C57BL6/J | Female | 103 | 105 | 107 | 98 | 1.84 | 143 | 146 | 144 | 140 | 1.39 |
| 78 | 50046787 | Mouse inbred strain C57BL6/J | Female | 125 | 122 | 128 | 126 | 3.01 | 149 | 147 | 155 | 144 | 4.22 |
| 79 | 50046785 | Mouse inbred strain C57BL6/J | Female | 144 | 148 | 143 | 141 | 2.65 | 173 | 177 | 169 | 174 | 4.00 |
| 80 | 50046773 | Mouse inbred strain C57BL6/J | Female | 55 | 58 | 53 | 55 | 2.50 | 71 | 73 | 69 | 72 | 2.01 |
| 81 | 50046777 | Mouse inbred strain C57BL6/J | Female | 74 | 71 | 77 | 73 | 3.01 | 135 | 138 | 129 | 139 | 4.62 |
| 82 | 50046786 | Mouse inbred strain C57BL6/J | Female | 225 | 223 | 221 | 232 | 2.17 | 258 | 259 | 256 | 260 | 1.58 |
| 83 | 50046778 | Mouse inbred strain C57BL6/J | Female | 116 | 119 | 115 | 113 | 2.14 | 166 | 164 | 168 | 165 | 2.01 |
| 84 | 50046770 | Mouse inbred strain C57BL6/J | Female | 118 | 120 | 114 | 121 | 3.10 | 139 | 137 | 135 | 144 | 1.84 |
| 85 | 50046762 | Mouse inbred strain C57BL6/J | Female | 122 | 123 | 119 | 124 | 2.08 | 168 | 167 | 164 | 173 | 2.08 |
| 86 | 50046788 | Mouse inbred strain C57BL6/J | Female | 100 | 101 | 104 | 96 | 1.95 | 141 | 139 | 143 | 140 | 2.01 |
| 87 | 50046775 | Mouse inbred strain C57BL6/J | Female | 181 | 176 | 186 | 182 | 5.00 | 209 | 206 | 212 | 208 | 3.01 |
| 88 | 50046781 | Mouse inbred strain C57BL6/J | Female | 288 | 293 | 281 | 291 | 6.05 | 302 | 305 | 306 | 296 | 1.90 |
| 89 | 50046792 | Mouse inbred strain C57BL6/J | Male | 54 | 51 | 59 | 55 | 4.04 | 85 | 89 | 86 | 87 | 2.08 |
| 90 | 50046797 | Mouse inbred strain C57BL6/J | Male | 83 | 89 | 90 | 87 | 3.79 | 134 | 138 | 132 | 135 | 3.06 |
| 91 | 50046801 | Mouse inbred strain C57BL6/J | Male | 99 | 91 | 90 | 93 | 4.93 | 137 | 130 | 139 | 135 | 4.73 |
| 92 | 50046806 | Mouse inbred strain C57BL6/J | Male | 177 | 182 | 185 | 181 | 4.04 | 228 | 226 | 222 | 225 | 3.06 |
| 93 | 50046807 | Mouse inbred strain C57BL6/J | Male | 208 | 193 | 195 | 199 | 8.14 | 223 | 229 | 230 | 227 | 3.79 |
| 94 | 50046809 | Mouse inbred strain C57BL6/J | Male | 161 | 163 | 157 | 160 | 3.06 | 219 | 225 | 218 | 221 | 3.79 |
| 95 | 50046814 | Mouse inbred strain C57BL6/J | Male | 172 | 173 | 169 | 171 | 2.08 | 220 | 227 | 216 | 221 | 5.57 |
| 96 | 50046815 | Mouse inbred strain C57BL6/J | Male | 176 | 179 | 168 | 174 | 5.69 | 221 | 222 | 228 | 224 | 3.79 |
| 97 | 50046816 | Mouse inbred strain C57BL6/J | Male | 117 | 120 | 116 | 118 | 2.08 | 161 | 160 | 164 | 162 | 2.08 |
| 98 | 50046820 | Mouse inbred strain C57BL6/J | Male | 108 | 103 | 107 | 106 | 2.65 | 142 | 148 | 146 | 145 | 3.06 |
| 99 | 50046763 | Mouse inbred strain C57BL6/J | Male | 204 | 201 | 196 | 200 | 4.04 | 229 | 229 | 224 | 227 | 2.89 |
| 100 | 50046764 | Mouse inbred strain C57BL6/J | Male | 193 | 206 | 200 | 200 | 6.51 | 235 | 229 | 230 | 231 | 3.21 |
| 101 | 50046765 | Mouse inbred strain C57BL6/J | Male | 227 | 229 | 220 | 225 | 4.73 | 261 | 259 | 253 | 258 | 4.16 |
| 102 | 50046766 | Mouse inbred strain C57BL6/J | Male | 150 | 153 | 154 | 152 | 2.08 | 203 | 207 | 206 | 205 | 2.08 |
| 103 | 50046767 | Mouse inbred strain C57BL6/J | Male | 158 | 152 | 156 | 155 | 3.06 | 211 | 209 | 207 | 209 | 2.00 |
| 104 | 50046769 | Mouse inbred strain C57BL6/J | Male | 158 | 157 | 155 | 157 | 1.53 | 213 | 220 | 214 | 216 | 3.79 |
| 105 | 50046772 | Mouse inbred strain C57BL6/J | Female | 165 | 158 | 157 | 160 | 4.36 | 219 | 223 | 222 | 221 | 2.08 |
| 106 | 50046774 | Mouse inbred strain C57BL6/J | Female | 80 | 83 | 81 | 81 | 1.53 | 134 | 128 | 135 | 132 | 3.79 |
| 107 | 50046776 | Mouse inbred strain C57BL6/J | Female | 101 | 98 | 102 | 100 | 2.08 | 145 | 143 | 136 | 141 | 4.73 |
| 108 | 50046779 | Mouse inbred strain C57BL6/J | Female | 97 | 103 | 99 | 100 | 3.06 | 139 | 147 | 142 | 143 | 4.04 |
| 109 | 50046780 | Mouse inbred strain C57BL6/J | Female | 117 | 121 | 115 | 118 | 3.06 | 167 | 163 | 157 | 162 | 5.03 |
| 110 | 50046784 | Mouse inbred strain C57BL6/J | Female | 118 | 123 | 115 | 119 | 4.04 | 163 | 160 | 171 | 165 | 5.69 |
| 111 | 50046790 | Mouse inbred strain C57BL6/J | Female | 128 | 124 | 129 | 127 | 2.65 | 172 | 169 | 177 | 173 | 4.04 |
| 112 |  |  |  |  |  |  | average SD: | 3.54 |  |  |  | average SD: | 3.15 |
| 113 |  |  |  |  |  |  |  |  |  |  |  |  |  |

**S2 Table.**
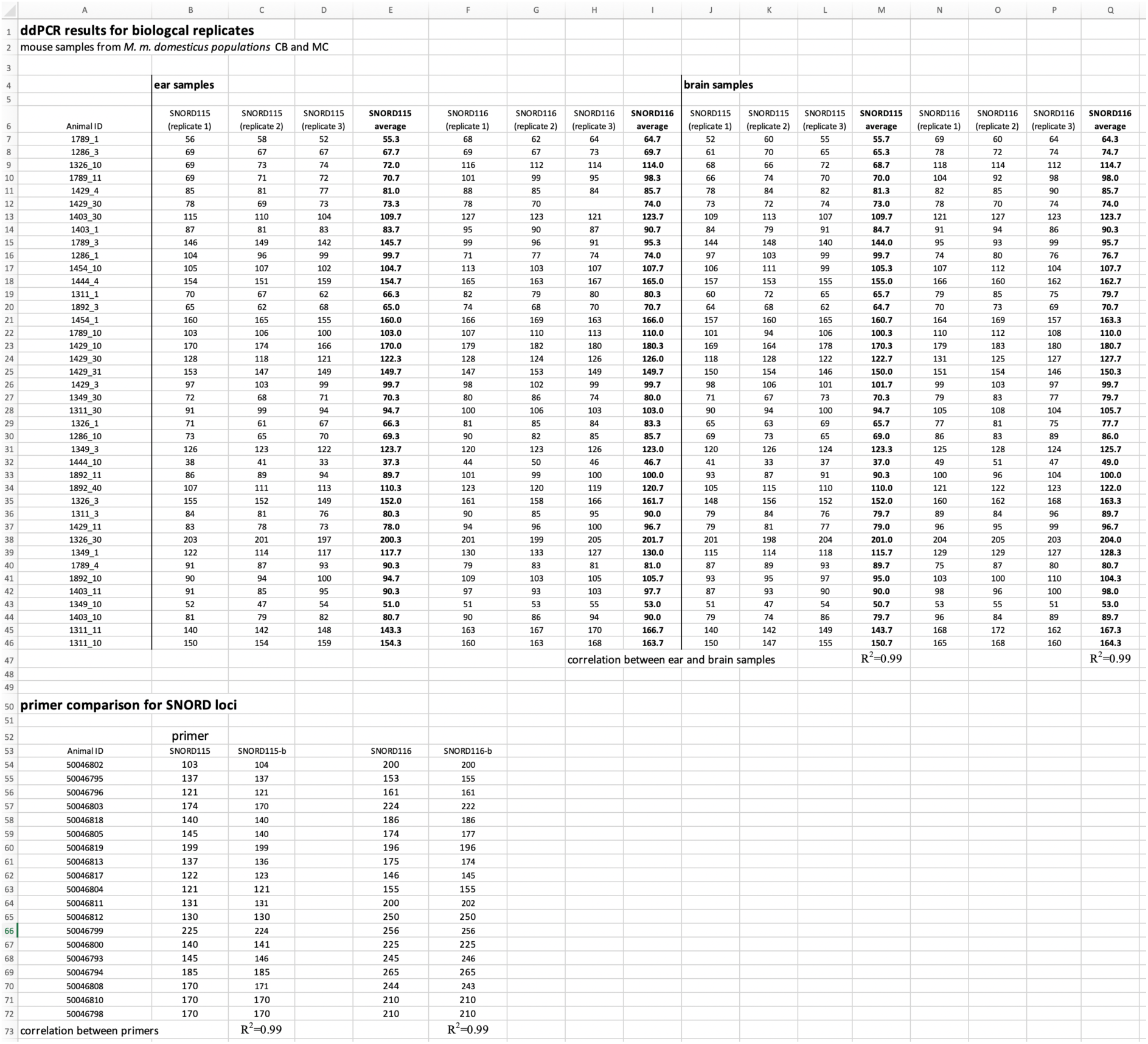

**S3 Table.**

|  | A | B | C | D | E | F | G | H | I | J | K | L | M | N | O | P | Q | R | S | T |
| --- | --- | --- | --- | --- | --- | --- | --- | --- | --- | --- | --- | --- | --- | --- | --- | --- | --- | --- | --- | --- |
| 1 | Lack of correlation with SNORD copy numbers to two other CNV loci (Sfi1 and CWC22) |  |  |  |  |  |  |  |  |  |  | Lack of behavioral correlation to two other CNV loci (Sfi and Cwc2) |  |  |  |  |  |  |  |  |
| 2 |  |  |  |  |  |  |  |  |  |  |  |  |  |  |  |  |  |  |  |  |
| 3 | Population CB |  |  |  |  |  |  |  |  |  |  | Population CB |  |  |  |  |  |  |  |  |
| 4 | Animal ID | Sfi copy number | SNORD115 copy number | SNORD116 copy number |  | Animal ID | Cwc22 copy number | SNORD115 copy number | SNORD116 copy number |  |  | Animal ID | Sfi copy number | Wall time | Dark arm time |  | Animal ID | Cwc22 copy number | Wall time | Dark arm time |
| 5 | 252980 | 29 | 95 | 132 |  | 252980 | 18 | 95 | 132 |  |  | 252980 | 29 | 86.68 | 150 |  | 252980 | 18 | 86.68 | 150 |
| 6 | 254434 | 33 | 195 | 224 |  | 254434 | 54 | 195 | 224 |  |  | 254434 | 33 | 243.84 | 115.32 |  | 254434 | 54 | 243.84 | 115.32 |
| 7 | 251505 | 69 | 177 | 206 |  | 251505 | 69 | 177 | 206 |  |  | 251505 | 69 | 124.48 | 101 |  | 251505 | 69 | 124.48 | 101 |
| 8 | 253612 | 53 | 130 | 155 |  | 253612 | 89 | 130 | 155 |  |  | 253612 | 53 | 94.64 | 158.4 |  | 253612 | 89 | 94.64 | 158.4 |
| 9 | 251736 | 44 | 166 | 197 |  | 251736 | 31 | 166 | 197 |  |  | 251736 | 44 | 165.76 | 113.64 |  | 251736 | 31 | 165.76 | 113.64 |
| 10 | 253911 | 98 | 145 | 171 |  | 253911 | 25 | 145 | 171 |  |  | 253911 | 98 | 193.2 | 123.16 |  | 253911 | 25 | 193.2 | 123.16 |
| 11 | 252388 | 25 | 131 | 167 |  | 252388 | 49 | 131 | 167 |  |  | 252388 | 25 | 169.6 | 106.28 |  | 252388 | 49 | 169.6 | 106.28 |
| 12 | 250737 | 18 | 152 | 180 |  | 250737 | 67 | 152 | 180 |  |  | 250737 | 18 | 157.4 | 163.08 |  | 250737 | 67 | 157.4 | 163.08 |
| 13 | 254089 | 19 | 226 | 243 |  | 254089 | 55 | 226 | 243 |  |  | 254089 | 19 | 215.28 | 158.48 |  | 254089 | 55 | 215.28 | 158.48 |
| 14 | 253366 | 47 | 205 | 193 |  | 253366 | 48 | 205 | 193 |  |  | 253366 | 47 | 198.12 | 124.56 |  | 253366 | 48 | 198.12 | 124.56 |
| 15 | 253721 | 42 | 212 | 245 |  | 253721 | 47 | 212 | 245 |  |  | 253721 | 42 | 208.24 | 146.4 |  | 253721 | 47 | 208.24 | 146.4 |
| 16 | 254154 | 46 | 218 | 241 |  | 254154 | 33 | 218 | 241 |  |  | 254154 | 46 | 209.08 | 151.84 |  | 254154 | 33 | 209.08 | 151.84 |
| 17 | 252264 | 27 | 220 | 245 |  | 252264 | 22 | 220 | 245 |  |  | 252264 | 27 | 136.92 | 165.1 |  | 252264 | 22 | 136.92 | 165.1 |
| 18 | 254321 | 28 | 243 | 255 |  | 254321 | 99 | 243 | 255 |  |  | 254321 | 28 | 229.44 | 137 |  | 254321 | 99 | 229.44 | 137 |
| 19 | 250148 | 42 | 233 | 273 |  | 250148 | 39 | 233 | 273 |  |  | 250148 | 42 | 208.12 | 213.68 |  | 250148 | 39 | 208.12 | 213.68 |
| 20 | 252408 | 37 | 263 | 294 |  | 252408 | 69 | 263 | 294 |  |  | 252408 | 37 | 181.04 | 144.56 |  | 252408 | 69 | 181.04 | 144.56 |
| 21 | 252909 | 85 | 202 | 228 |  | 252909 | 71 | 202 | 228 |  |  | 252909 | 85 | 131 | 155.44 |  | 252909 | 71 | 131 | 155.44 |
| 22 | 252396 | 71 | 273 | 322 |  | 252396 | 74 | 273 | 322 |  |  | 252396 | 71 | 231.64 | 183.76 |  | 252396 | 74 | 231.64 | 183.76 |
| 23 | 254175 | 50 | 290 | 324 |  | 254175 | 39 | 290 | 324 |  |  | 254175 | 50 | 242 | 221.96 |  | 254175 | 39 | 242 | 221.96 |
| 24 | 250230 | 56 | 290 | 325 |  | 250230 | 64 | 290 | 325 |  |  | 250230 | 56 | 245.08 | 202.96 |  | 250230 | 64 | 245.08 | 202.96 |
| 25 | 253670 | 46 | 230 | 231 |  | 253670 | 35 | 230 | 231 |  |  | 253670 | 46 | 230.88 | 248.1 |  | 253670 | 35 | 230.88 | 248.1 |
| 26 | 251786 | 36 | 146 | 190 |  | 251786 | 52 | 146 | 190 |  |  | 251786 | 36 | 183.04 | 122.2 |  | 251786 | 52 | 183.04 | 122.2 |
| 27 |  |  |  |  |  |  |  |  |  |  |  |  |  |  |  |  |  |  |  |  |
| 28 | R<sup>2</sup> |  | 0.007 | 0.01 |  |  |  | 0.04 | 0.04 |  |  | R<sup>2</sup> |  | 0.0006 | 0.0004 |  |  |  | 0.001 | 0.01 |
| 29 | P-value |  | 0.71 | 0.65 |  |  |  | 0.35 | 0.36 |  |  | P-value |  | 0.92 | 0.93 |  |  |  | 0.88 | 0.65 |
| 30 |  |  |  |  |  |  |  |  |  |  |  |  |  |  |  |  |  |  |  |  |
| 31 |  |  |  |  |  |  |  |  |  |  |  |  |  |  |  |  |  |  |  |  |
| 32 | Population MC |  |  |  |  |  |  |  |  |  |  | Population MC |  |  |  |  |  |  |  |  |
| 33 | 250687 | 85 | 137 | 137 |  |  |  |  |  |  |  | 250687 | 85 | 193.92 | 100.04 |  |  |  |  |  |
| 34 | 254940 | 121 | 108 | 86 |  |  |  |  |  |  |  | 254940 | 121 | 187.6 | 106.8 |  |  |  |  |  |
| 35 | 250556 | 49 | 115 | 110 |  |  |  |  |  |  |  | 250556 | 49 | 199.44 | 223.84 |  |  |  |  |  |
| 36 | 250622 | 63 | 127 | 116 |  |  |  |  |  |  |  | 250622 | 63 | 195.24 | 119.68 |  |  |  |  |  |
| 37 | 253429 | 75 | 133 | 127 |  |  |  |  |  |  |  | 253429 | 75 | 195.76 | 186.36 |  |  |  |  |  |
| 38 | 254298 | 58 | 139 | 160 |  |  |  |  |  |  |  | 254298 | 58 | 215.64 | 174.88 |  |  |  |  |  |
| 39 | 251209 | 79 | 155 | 190 |  |  |  |  |  |  |  | 251209 | 79 | 276.84 | 195 |  |  |  |  |  |
| 40 | 254488 | 39 | 157 | 200 |  |  |  |  |  |  |  | 254488 | 39 | 292.52 | 162.88 |  |  |  |  |  |
| 41 | 253218 | 61 | 160 | 213 |  |  |  |  |  |  |  | 253218 | 61 | 237.24 | 176.12 |  |  |  |  |  |
| 42 | 250481 | 52 | 169 | 185 |  |  |  |  |  |  |  | 250481 | 52 | 216.92 | 210.44 |  |  |  |  |  |
| 43 | 250400 | 112 | 175 | 220 |  |  |  |  |  |  |  | 250400 | 112 | 251.88 | 136.8 |  |  |  |  |  |
| 44 | 250061 | 55 | 160 | 170 |  |  |  |  |  |  |  | 250061 | 55 | 230.04 | 150.04 |  |  |  |  |  |
| 45 | 252446 | 49 | 156 | 162 |  |  |  |  |  |  |  | 252446 | 49 | 239.64 | 162.32 |  |  |  |  |  |
| 46 | 254418 | 76 | 160 | 185 |  |  |  |  |  |  |  | 254418 | 76 | 236 | 217.36 |  |  |  |  |  |
| 47 | 251023 | 69 | 190 | 205 |  |  |  |  |  |  |  | 251023 | 69 | 236.36 | 140.76 |  |  |  |  |  |
| 48 | 250053 | 36 | 221 | 241 |  |  |  |  |  |  |  | 250053 | 36 | 257.96 | 184.44 |  |  |  |  |  |
| 49 | 254918 | 95 | 229 | 249 |  |  |  |  |  |  |  | 254918 | 95 | 244 | 235 |  |  |  |  |  |
| 50 | 251016 | 81 | 168 | 172 |  |  |  |  |  |  |  | 251016 | 81 | 236.36 | 233.76 |  |  |  |  |  |
| 51 | 251249 | 18 | 237 | 257 |  |  |  |  |  |  |  | 251249 | 18 | 255.64 | 175.12 |  |  |  |  |  |
| 52 | 253816 | 67 | 246 | 267 |  |  |  |  |  |  |  | 253816 | 67 | 261.96 | 207 |  |  |  |  |  |
| 53 | 250545 | 69 | 220 | 290 |  |  |  |  |  |  |  | 250545 | 69 | 286.52 | 263.3 |  |  |  |  |  |
| 54 | 252169 | 114 | 172 | 169 |  |  |  |  |  |  |  | 252169 | 114 | 185.08 | 202 |  |  |  |  |  |
| 55 | 251664 | 99 | 185 | 220 |  |  |  |  |  |  |  | 251664 | 99 | 262.88 | 260.1 |  |  |  |  |  |
| 56 |  |  |  |  |  |  |  |  |  |  |  |  |  |  |  |  |  |  |  |  |
| 57 | R<sup>2</sup> |  | 0.04 | 0.05 |  |  |  |  |  |  |  | R<sup>2</sup> |  | 0.09 | 0.0009 |  |  |  |  |  |
| 58 | P-value |  | 0.34 | 0.32 |  |  |  |  |  |  |  | P-value |  | 0.16 | 0.89 |  |  |  |  |  |
| 59 |  |  |  |  |  |  |  |  |  |  |  |  |  |  |  |  |  |  |  |  |

**S4 Table.**

|  | A | B | C | D | E | F | G | H | I |
| --- | --- | --- | --- | --- | --- | --- | --- | --- | --- |
| 1 | relative RNA expression in the brain, based on ddPCR scores |  |  |  |  |  |  |  |  |
| 2 |  |  |  |  |  |  |  |  |  |
| 3 | MouseID | Population | sex | SNORD 115 copy number | SNORD 115 expression | Htr2c ExonV expression | SNORD116 copy number | SNORD 116 expression | ANKRD11 exon 10 expression |
| 22 | 251664 | MC | Male | 185 | 211 | 220 | 220 | 138 | 140 |
| 23 |  | correlation of copy number with SNORD expression |  |  | R<sup>2</sup>=0.93 |  |  | R<sup>2</sup>=0.53 |  |
| 24 |  | correlation of SNORD expression with target exon expression |  |  |  | R<sup>2</sup>=0.65 |  |  | R<sup>2</sup>=0.72 |
| 25 |  |  |  |  |  |  |  |  |  |
| 26 | MouseID | Population | sex | SNORD 115 copy number | SNORD 115 expression | Htr2c ExonV expression | SNORD116 copy number | SNORD 116 expression | ANKRD11 exon 10 expression |
| 27 | 252980 | CB | Male | 95 | 132 | 267 | 131 | 173 | 227 |
| 28 | 251736 | CB | Male | 166 | 146 | 281 | 197 | 186 | 244 |
| 29 | 254089 | CB | Male | 226 | 188 | 255 | 243 | 227 | 363 |
| 30 | 252264 | CB | Male | 220 | 167 | 302 | 245 | 239 | 387 |
| 31 | 250148 | CB | Male | 233 | 211 | 345 | 270 | 275 | 417 |
| 32 | 252396 | CB | Male | 275 | 298 | 393 | 320 | 339 | 479 |
| 33 | 250230 | CB | Male | 291 | 281 | 416 | 325 | 362 | 460 |
| 34 | 251786 | CB | Male | 146 | 227 | 405 | 190 | 287 | 339 |
| 35 | 250687 | MC | Male | 137 | 142 | 171 | 137 | 66 | 83 |
| 36 | 254940 | MC | Male | 108 | 84 | 151 | 86 | 112 | 75 |
| 37 | 254298 | MC | Male | 139 | 115 | 153 | 160 | 77 | 47 |
| 38 | 253218 | MC | Male | 160 | 157 | 193 | 213 | 89 | 96 |
| 39 | 252446 | MC | Male | 156 | 164 | 207 | 160 | 116 | 111 |
| 40 | 254918 | MC | Male | 221 | 230 | 227 | 249 | 156 | 129 |
| 41 | 250545 | MC | Male | 220 | 232 | 235 | 290 | 187 | 149 |
| 42 | 251664 | MC | Male | 185 | 211 | 220 | 220 | 138 | 140 |
| 43 |  | correlation of copy number with SNORD expression |  |  | R<sup>2</sup>=0.71 |  |  | R<sup>2</sup>=0.53 |  |
| 44 |  | correlation of SNORD expression with target exon expression |  |  |  | R<sup>2</sup>=0.56 |  |  | R<sup>2</sup>=0.91 |
| 45 |  |  |  |  |  |  |  |  |  |

**S5 Table.**
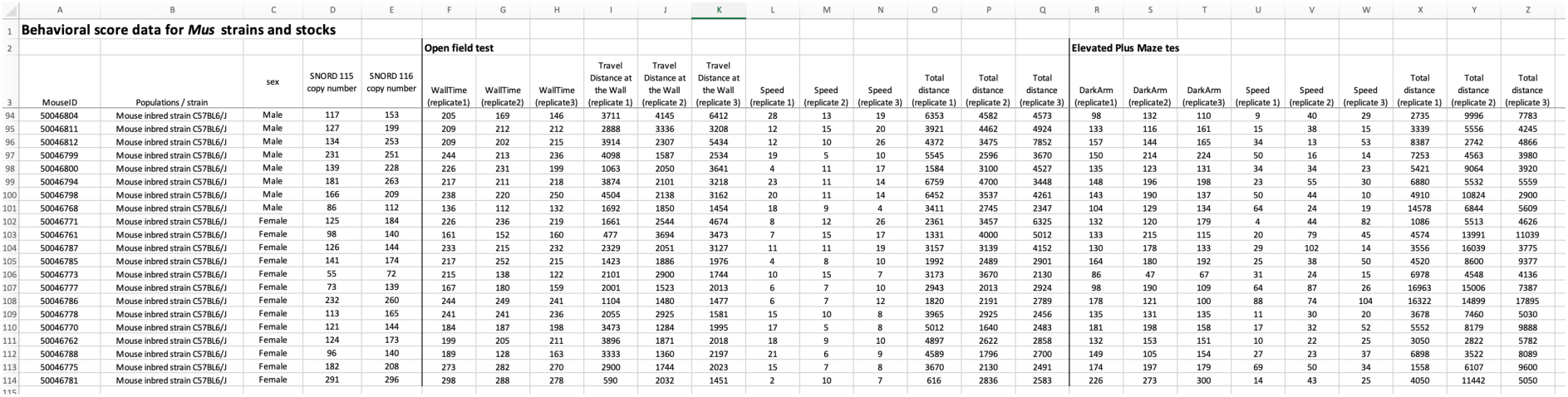

**S6 Table.**
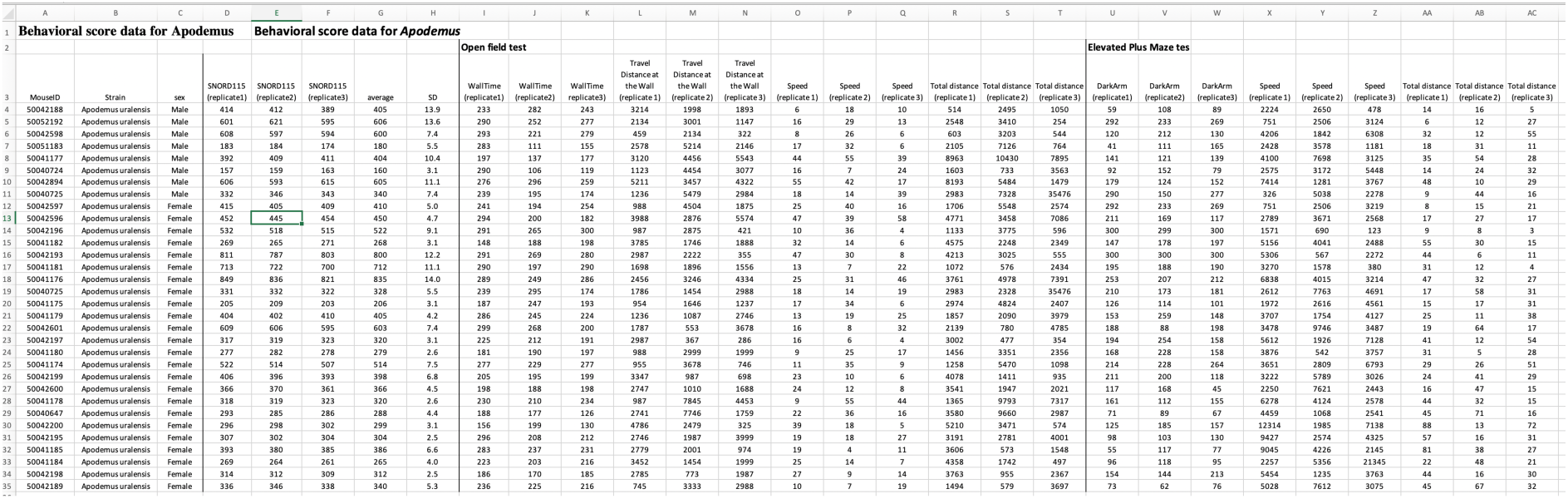

**S7 Table.**
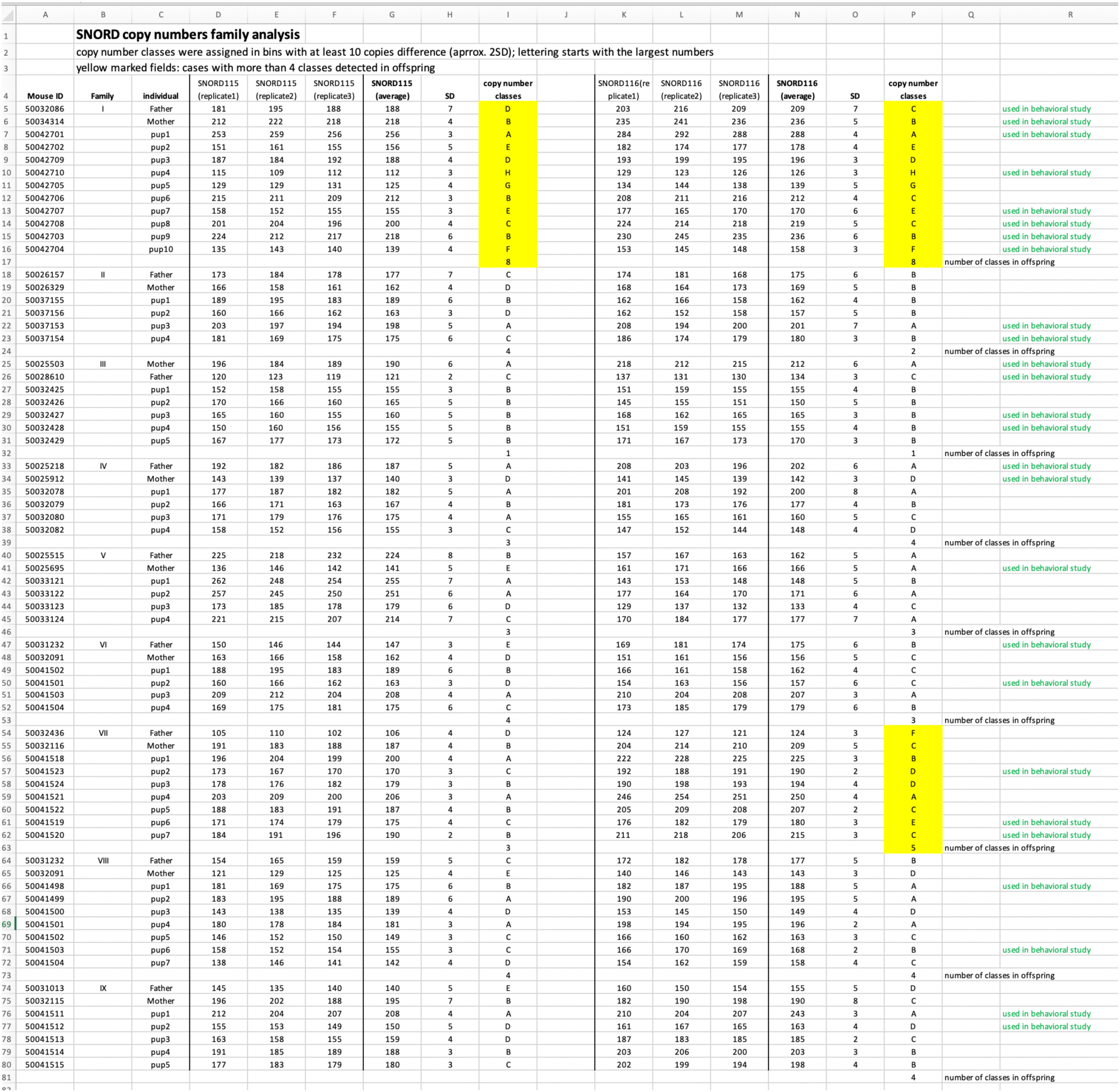

**S8 Table.**
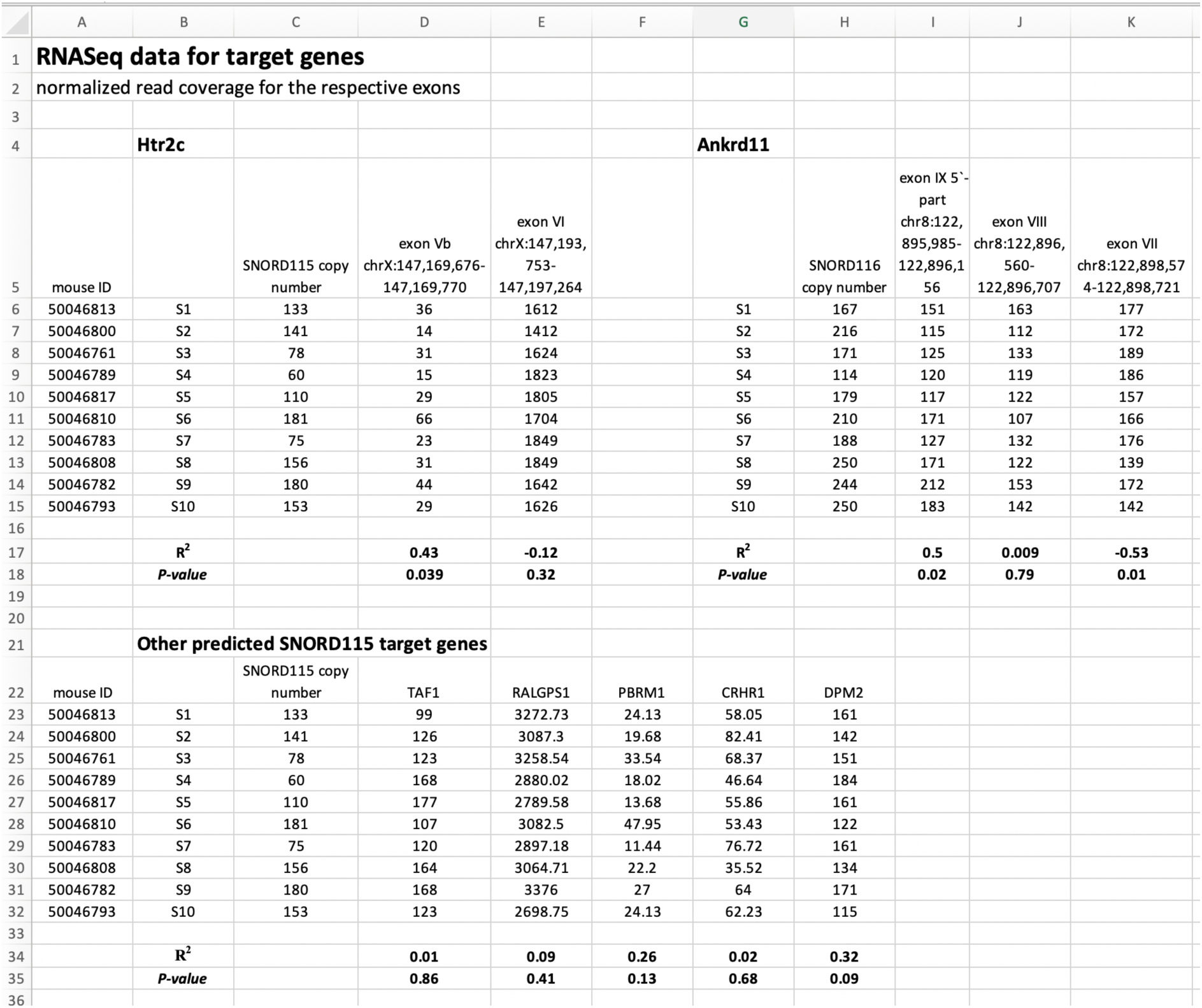

|  | A | B | C | D | E | F | G | H | I | J | K |
| --- | --- | --- | --- | --- | --- | --- | --- | --- | --- | --- | --- |
| 1 | **RNASeq data for target genes** |  |  |  |  |  |  |  |  |  |  |
| 2 | normalized read coverage for the respective exons |  |  |  |  |  |  |  |  |  |  |
| 3 |  |  |  |  |  |  |  |  |  |  |  |
| 4 |  | **Htr2c** |  |  |  |  | **Ankrd11** |  |  |  |  |
| 5 | mouse ID |  | SNORD115 copy number | exon Vb<br>chrX:147,169,676-147,169,770 | exon VI<br>chrX:147,193,753-147,197,264 |  |  | SNORD116 copy number | exon IX 5'-part<br>chr8:122,895,985-122,896,156 | exon VIII<br>chr8:122,896,560-122,896,707 | exon VII<br>chr8:122,898,574-122,898,721 |
| 6 | 50046813 | S1 | 133 | 36 | 1612 |  | S1 | 167 | 151 | 163 | 177 |
| 7 | 50046800 | S2 | 141 | 14 | 1412 |  | S2 | 216 | 115 | 112 | 172 |
| 8 | 50046761 | S3 | 78 | 31 | 1624 |  | S3 | 171 | 125 | 133 | 189 |
| 9 | 50046789 | S4 | 60 | 15 | 1823 |  | S4 | 114 | 120 | 119 | 186 |
| 10 | 50046817 | S5 | 110 | 29 | 1805 |  | S5 | 179 | 117 | 122 | 157 |
| 11 | 50046810 | S6 | 181 | 66 | 1704 |  | S6 | 210 | 171 | 107 | 166 |
| 12 | 50046783 | S7 | 75 | 23 | 1849 |  | S7 | 188 | 127 | 132 | 176 |
| 13 | 50046808 | S8 | 156 | 31 | 1849 |  | S8 | 250 | 171 | 122 | 139 |
| 14 | 50046782 | S9 | 180 | 44 | 1642 |  | S9 | 244 | 212 | 153 | 172 |
| 15 | 50046793 | S10 | 153 | 29 | 1626 |  | S10 | 250 | 183 | 142 | 142 |
| 16 |  |  |  |  |  |  |  |  |  |  |  |
| 17 |  | **R<sup>2</sup>** |  | **0.43** | **-0.12** |  | **R<sup>2</sup>** |  | **0.5** | **0.009** | **-0.53** |
| 18 |  | **P-value** |  | **0.039** | **0.32** |  | **P-value** |  | **0.02** | **0.79** | **0.01** |
| 19 |  |  |  |  |  |  |  |  |  |  |  |
| 20 |  |  |  |  |  |  |  |  |  |  |  |
| 21 | **Other predicted SNORD115 target genes** |  |  |  |  |  |  |  |  |  |  |
| 22 | mouse ID |  | SNORD115 copy number | TAF1 | RALGPS1 | PBRM1 | CRHR1 | DPM2 |  |  |  |
| 23 | 50046813 | S1 | 133 | 99 | 3272.73 | 24.13 | 58.05 | 161 |  |  |  |
| 24 | 50046800 | S2 | 141 | 126 | 3087.3 | 19.68 | 82.41 | 142 |  |  |  |
| 25 | 50046761 | S3 | 78 | 123 | 3258.54 | 33.54 | 68.37 | 151 |  |  |  |
| 26 | 50046789 | S4 | 60 | 168 | 2880.02 | 18.02 | 46.64 | 184 |  |  |  |
| 27 | 50046817 | S5 | 110 | 177 | 2789.58 | 13.68 | 55.86 | 161 |  |  |  |
| 28 | 50046810 | S6 | 181 | 107 | 3082.5 | 47.95 | 53.43 | 122 |  |  |  |
| 29 | 50046783 | S7 | 75 | 120 | 2897.18 | 11.44 | 76.72 | 161 |  |  |  |
| 30 | 50046808 | S8 | 156 | 164 | 3064.71 | 22.2 | 35.52 | 134 |  |  |  |
| 31 | 50046782 | S9 | 180 | 168 | 3376 | 27 | 64 | 171 |  |  |  |
| 32 | 50046793 | S10 | 153 | 123 | 2698.75 | 24.13 | 62.23 | 115 |  |  |  |
| 33 |  |  |  |  |  |  |  |  |  |  |  |
| 34 |  | **R<sup>2</sup>** |  | **0.01** | **0.09** | **0.26** | **0.02** | **0.32** |  |  |  |
| 35 |  | **P-value** |  | **0.86** | **0.41** | **0.13** | **0.68** | **0.09** |  |  |  |
| 36 |  |  |  |  |  |  |  |  |  |  |  |

**S9 Table.**
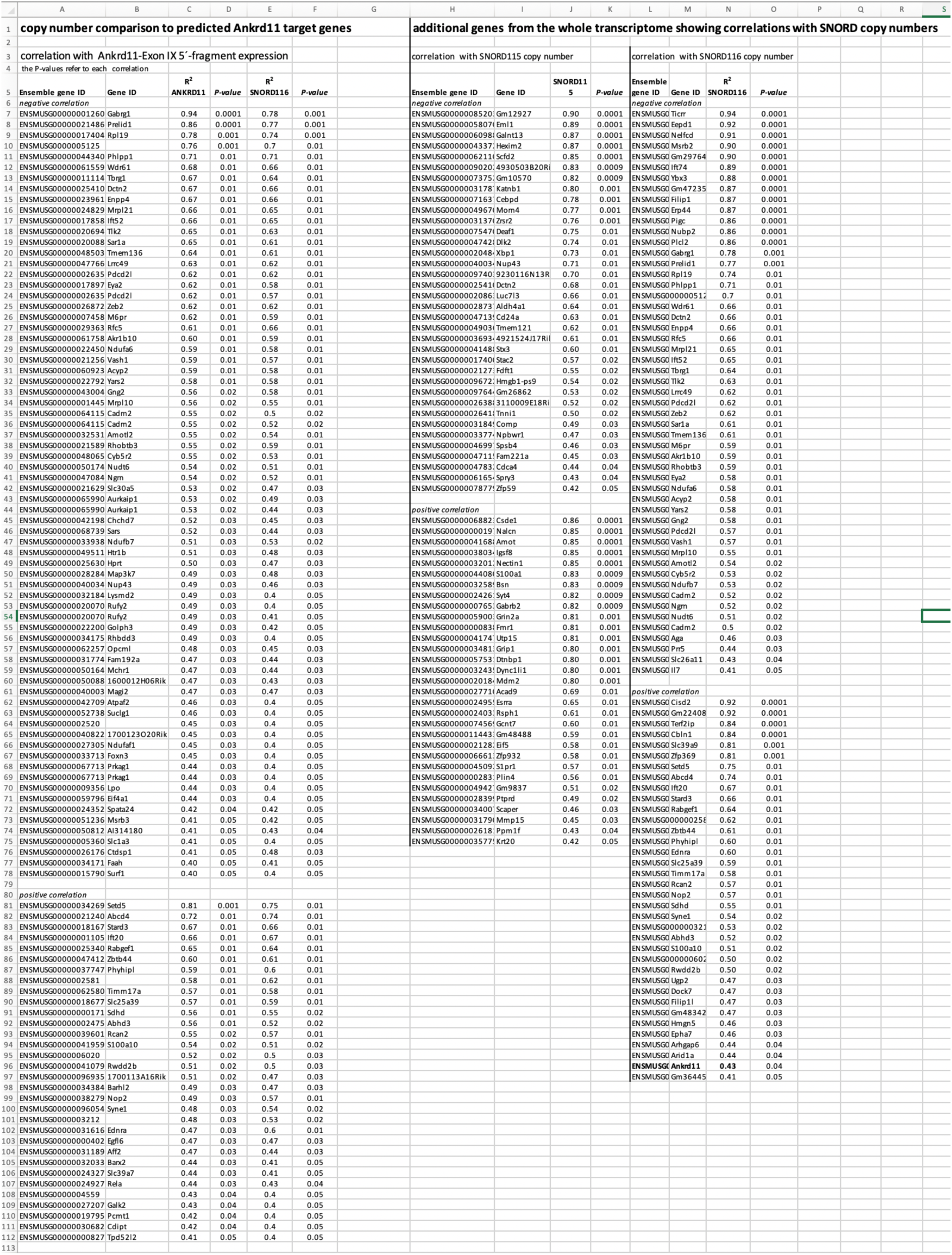

**S10 Table.**

|  | A | B | C | D | E | F | G | H | I |
| --- | --- | --- | --- | --- | --- | --- | --- | --- | --- |
| 1 | Behavioral score data for *Cavea* |  |  |  |  |  |  |  |  |
| 2 | animal ID | sex | SNORD116<br>(replicate 1) | SNORD116<br>(replicate 2) | SNORD116<br>(replicate 3) | average | SD | anxiety score in the<br>Open Field test<br>(distance run in cm) | anxiety<br>scoreStruggle<br>Docility (seconds) |
| 3 | 72A-C0B7 | Male | 10 | 12 | 11 | 11 | 1.0 | 6560 | 14 |
| 4 | 74F-E335 | Male | 9 | 10 | 12 | 10 | 1.5 | 3168 | 13 |
| 5 | 74F-B698 | Male | 12 | 13 | 10 | 12 | 1.5 | 5417 | 13 |
| 6 | 72A-BA76 | Male | 13 | 11 | 12 | 12 | 1.0 | 322 | 11 |
| 7 | 72A-8511 | Male | 7 | 8 | 10 | 8 | 1.5 | 8034 | 20 |
| 8 | 75B-460E | Male | 11 | 10 | 8 | 10 | 1.5 | 4896 | 15 |
| 9 | 72A-808E | Male | 8 | 6 | 10 | 8 | 2.0 | 21942 | 16 |
| 10 | 72A-ABED | Male | 13 | 12 | 11 | 12 | 1.0 | 3942 | 10 |
| 11 | 75B-72C0 | Male | 9 | 6 | 10 | 8 | 2.1 | 5284 | 19 |
| 12 | 3433 | Female | 13 | 10 | 9 | 11 | 2.1 | 5152 | 14 |
| 13 | 4246 | Female | 15 | 16 | 12 | 14 | 2.1 | 356 | 2 |
| 14 | D080 | Female | 7 | 9 | 8 | 8 | 1.0 | 5783 | 22 |
| 15 | 4F77 | Female | 9 | 10 | 13 | 11 | 2.1 | 1585 | 1 |
| 16 | 7CB4 | Female | 9 | 13 | 10 | 11 | 2.1 | 997 | 16 |
| 17 | 8326 | Female | 9 | 11 | 15 | 12 | 3.1 | 0 | 17 |
| 18 | 36C2 | Female | 11 | 9 | 8 | 9 | 1.5 | 4987 | 10 |
| 19 | 8341 | Female | 10 | 12 | 9 | 10 | 1.5 | 1763 | 16 |
| 20 | 390D | Female | 6 | 8 | 8 | 7 | 1.2 | 4212 | 24 |
| 21 | 0FCF | Female | 8 | 8 | 10 | 9 | 1.2 | 1873 | 12 |

**S11 Table.**
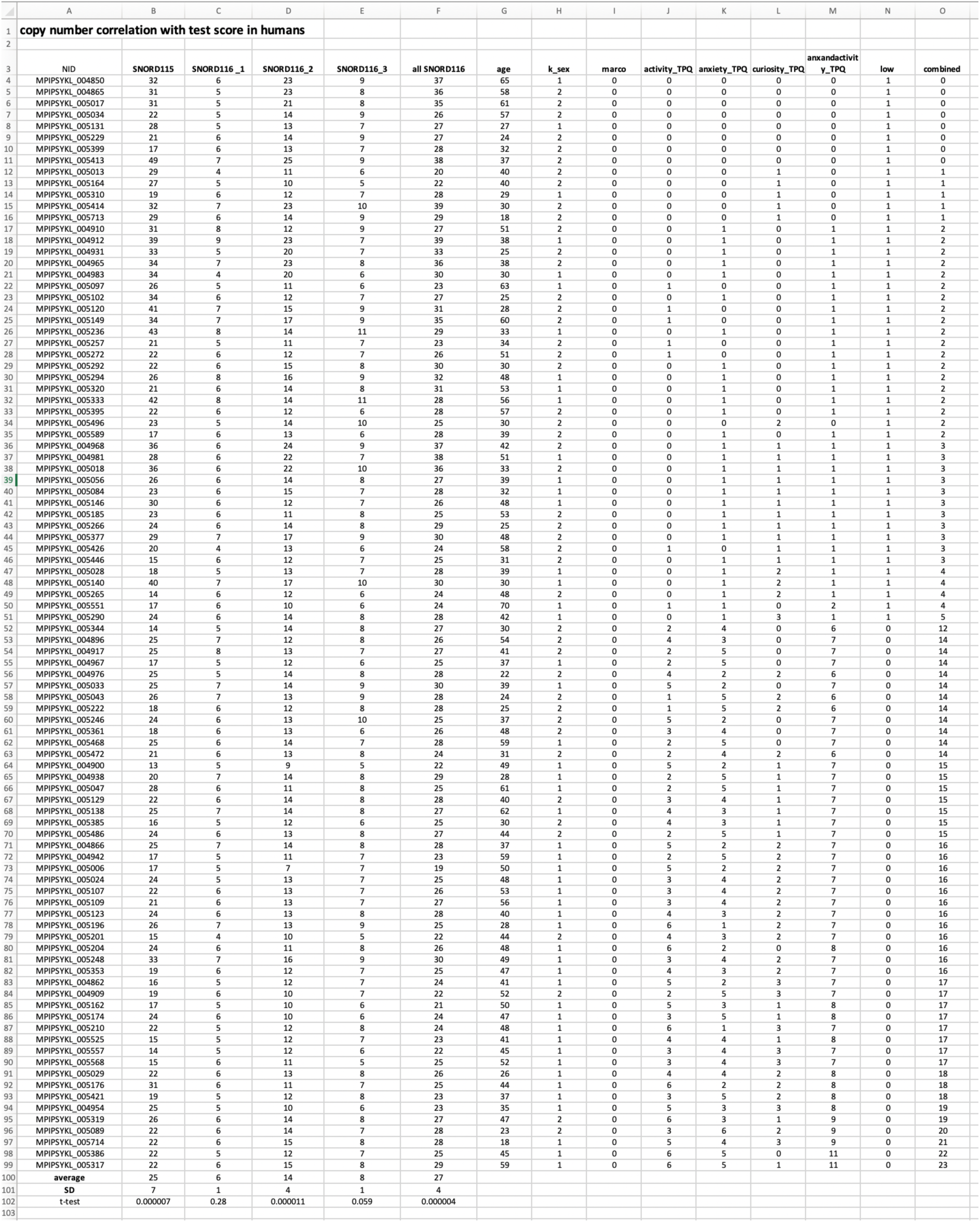

